# Mechanisms of transcriptional regulation in *Anopheles gambiae* revealed by allele specific expression

**DOI:** 10.1101/2023.11.22.568226

**Authors:** Naomi A. Dyer, Eric R. Lucas, Sanjay C. Nagi, Daniel P. McDermott, Jon H. Brenas, Alistair Miles, Chris S. Clarkson, Henry D. Mawejje, Craig S. Wilding, Marc S. Halfon, Hasiba Asma, Eva Heinz, Martin J. Donnelly

## Abstract

Malaria control relies on insecticides targeting the mosquito vector, but this is increasingly compromised by insecticide resistance, which can be achieved by elevated expression of detoxifying enzymes that metabolize the insecticide. In diploid organisms, gene expression is regulated both in *cis*, by regulatory sequences on the same chromosome, and by *trans* acting factors, affecting both alleles equally. Differing levels of transcription can be caused by mutations in *cis*-regulatory modules (CRM), but few of these have been identified in mosquitoes. We crossed bendiocarb resistant and susceptible *Anopheles gambiae* strains to identify *cis*-regulated genes that might be responsible for the resistant phenotype using RNAseq, and *cis*-regulatory module sequences controlling gene expression in insecticide resistance relevant tissues were predicted using machine learning. We found 115 genes showing allele specific expression in hybrids of insecticide susceptible and resistant strains, suggesting *cis* regulation is an important mechanism of gene expression regulation in *Anopheles gambiae*. The genes showing allele specific expression included a higher proportion of *Anopheles* specific genes on average younger than genes those with balanced allelic expression.

**Author Summary:** The evolution of insecticide resistance, including resistance that is due to changes in the expression levels of certain resistance associated genes is threatening progress in malaria control. We investigated how the expression of genes in the malaria vector *Anopheles gambiae* is controlled, by implementing a method for the first time in this species. Each mosquito inherits a set of chromosomes from both parents, so has a maternal and paternal copy of most genes. When a gene is expressed, the DNA encoding that gene is transcribed into messenger RNA. This process is controlled by the cellular environment and by other DNA sequences on the same chromosome as each gene. We crossed mosquitoes from insecticide resistant and susceptible strains to equalize the cellular environment and then measured the levels of messenger RNA from both gene copies. 115 genes showed consistently different messenger RNA levels between gene copies in most crosses, suggesting these genes are regulated by factors on the same chromosome. There were relatively more Anopheles specific genes with imbalanced expression. Using machine learning we identified DNA sequences that may be responsible for controlling gene expression in mosquito tissues; several of these sequences were close to genes with imbalanced expression.

## Introduction

Malaria prevalence in Sub-Saharan Africa has reduced by 50% since 2000, primarily due to insecticide-based control of mosquito vectors [1]. Recently, progress has stagnated [2], partly due to increasing levels of resistance against insecticides in mosquito populations [3]. *Anopheles gambiae* is one of the dominant malaria vectors in Sub-Saharan Africa, the primary vector across most of Uganda [4] and the vector for which the largest resource of genome data is available, with 7275 genomes sequenced [5–8]. A common cause of insecticide resistance is increased degradation of insecticides (termed metabolic resistance) [9] with overexpression of insecticide-metabolizing P450s repeatedly implicated [10–12]. This can be caused by mutations in *cis*-regulatory regions regulating expression of metabolic resistance genes. In diploid organisms, such mutations only affect expression of the allele of the gene located on the same chromosome. Although some *trans* factors involved in metabolic resistance gene regulation in *Anopheles* are known [13, 14], few studies have identified genetic variation causing metabolic resistance [15–18]. The multiallelic nature of metabolic insecticide resistance which can involve different mutations affecting the same gene in different populations, as well as the involvement of multiple genes, makes marker identification challenging as it limits the power of association studies unless very large sample sizes are used [15]. Despite the primary role of gene overexpression in metabolic resistance, only one *cis*-regulatory variant for resistance-linked differential expression has been identified in *Anopheles gambiae* [19], and markers for such variants are therefore absent in the current genetic marker panel for resistance [20]. Copy number variants (CNV) have been observed in *Anopheles gambiae* metabolic resistance gene clusters [21]. For example, in *Anopheles coluzzi* copy number of *Cyp6AA1* is associated with deltamethrin resistance [22]. The relative contribution of CNV and *cis*-regulation on *Anopheles* gene expression has not yet been determined.

Uganda sees a high burden of malaria; comprising 7.8% of all global cases in 2021 [23]. To address this public health burden, the insecticide bendiocarb has been used for indoor residual spraying (IRS) to complement the distribution of long-lasting insecticidal nets (LLIN). Some resistance to bendiocarb in *An*. *gambiae* was observed in Nagongera (southeast Uganda) and Kihihi (southwest Uganda), with 83% and 70% mortality, respectively, to WHO bioassays with a diagnostic dose of 0.1% bendiocarb in 2014 prior to the IRS [24]. The IRS campaign starting in December 2014 succeeded in reducing human biting rate, cases of malaria and test positivity rate in Nagongera [24], but the potential for further increases in resistance puts the long-term usefulness of bendiocarb into question.

Mosquitoes collected from Nagongera in 2014, which have moderate resistance to bendiocarb, showed significant differential expression of many genes compared to the susceptible Kisumu strain, including salivary gland protein encoding *D7r2* and *D7r4* genes as well as the detoxification associated genes *Gstd3* and *Cyp6m2* [25]. Expression of *D7r4* was associated with a single-nucleotide polymorphism (SNP) in a non-coding transcript downstream of the D7 cassette [25].

In diploid organisms, allele specific expression (ASE) provides strong evidence that genes may be under differential *cis*-regulatory control [26, 27]. Using a method that has been applied to a variety of taxa [26, 28–32] but not mosquitoes, we describe the identification of genes showing ASE in *An. gambiae* which potentially confer metabolic resistance. The challenges of applying this method that arise from mosquito biology and genome structure are discussed. In complementary work we predicted the sequences of some of the *cis* regulatory modules that may underlie the expression of genes involved in insecticide resistance using machine learning. Predictions included potential CRMs proximal to the genes showing ASE and genes that show consistent differential expression patterns in multiple resistant *Anopheles* strains, providing a starting point for future investigations into CRM variants during the evolution of insecticide resistance.

## Materials and Methods

### Strains and Crosses

Resting female mosquitoes were collected in Siwa Village, Nagongera, Tororo District in Eastern Uganda (0°46’12.0"N, 34°01’34.0"E) in March 2013 and allowed to lay eggs. 65 *Anopheles gambiae* (as ascertained by species ID PCR [33]) egg batches laid by the collected females were reared to larvae and combined to establish the Nagongera colony. All the colony founding mothers were screened for a known bendiocarb target site resistance mutation G119S in the *Ace1* gene using the TaqMan assay described in [34]. No resistance associated variants were observed and full sequencing of the *Ace1* gene from six colony mosquitoes that survived Bendiocarb exposure did not reveal any other potential target site resistance mutations. The colony was assayed using WHO tube assays [35] at F2 and was found to be highly resistant to DDT (at diagnostic dose 4%) and deltamethrin (at diagnostic dose 0.05%), intermediate resistance to the carbamate insecticide bendiocarb (at diagnostic dose 0.1%). Bioassays of Nagongera colony adults using the WHO tube assay [35, 36] with preexposure to 4% piperonyl butoxide (which inhibits P450 mediated metabolism) for one hour prior to bendiocarb exposure increased mortality compared to bendiocarb exposure alone, implying that bendiocarb resistance is metabolic rather than target site mediated. A schematic of the experimental design is shown in figure 1. In brief, reciprocal crosses between Nagongera mosquitoes and an inbred insecticide susceptible colony (origin Kisumu, Kenya, 1975) were performed in cages as described previously [6]. Reciprocal crosses each involved a total of 13 males and multiple females; since *Anopheles gambiae* are swarm maters, multiple males are required to induce mating. Following mating, females were transferred to individual cups for egg laying. F1 progeny of three females from each reciprocal cross were raised to adults under standard insectary conditions of 12 hours light 12 hours dark cycle, 26°C +/-2°C and 70% relative humidity, and fed on 10% sucrose solution (Table 1, Figure 1).

**Figure 1.**
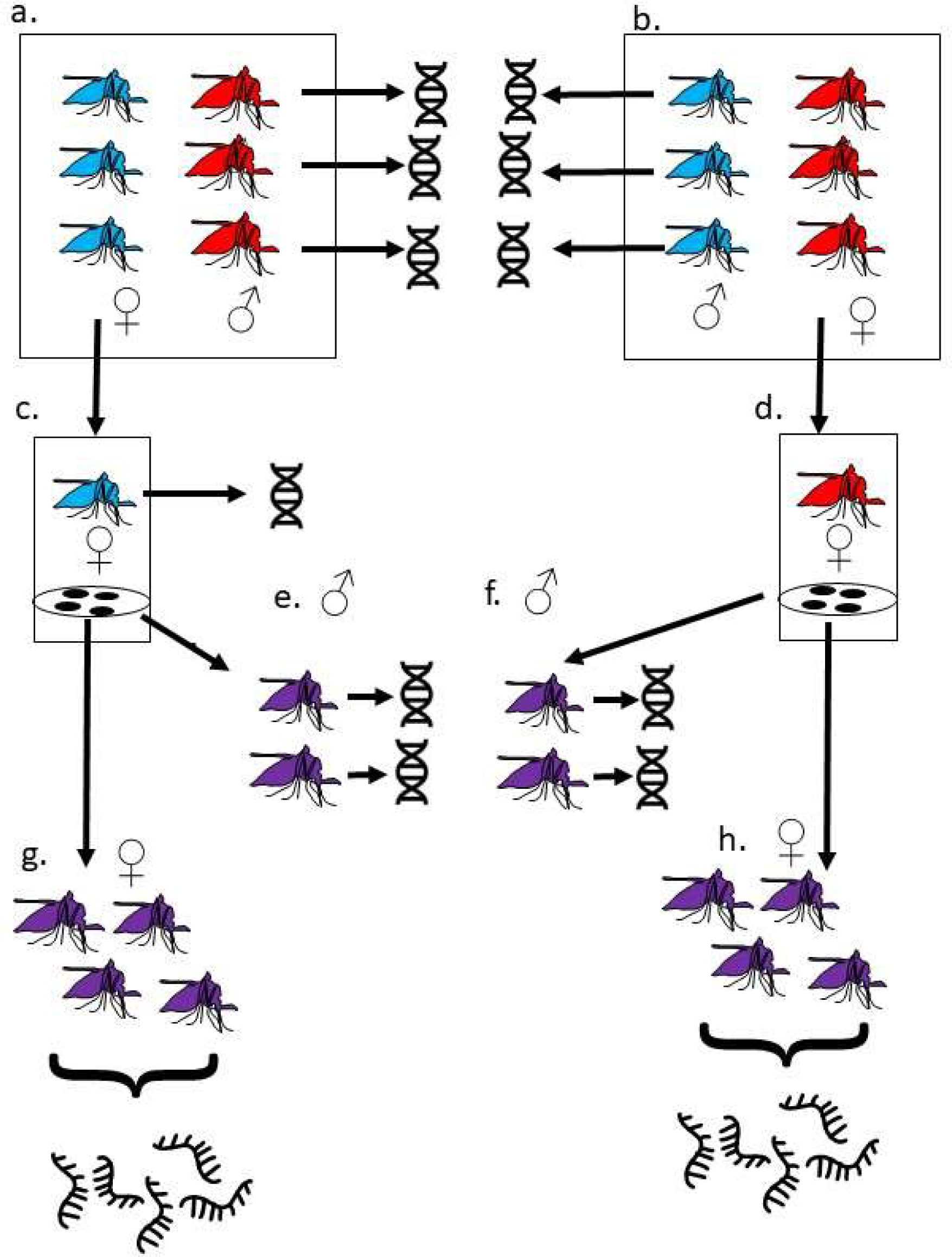
Crossing, DNA and RNA extraction schema. a. 13 Kisumu females (blue) were crossed to 13 Nagongera males (red). Females mate only once. Following mating, genomic DNA from Nagongera males was extracted and sequenced. b. In the reciprocal cross, 13 Nagongera females (red) were crossed to 13 Kisumu males (blue). Following mating, genomic DNA was extracted and sequenced from Kisumu males. c. Individual mated Kisumu females were transferred to laying cups. Following egg laying, genomic DNA was extracted and sequenced from three of these d. Individual mated Nagongera females were transferred to laying cups. Following egg laying, genomic DNA was extracted and sequenced from three of these. e. The F1 progeny (purple) from the three Kisumu mothers sequenced at step c. were raised to adulthood. Genomic DNA was extracted and sequenced from individual males f. The F1 progeny (purple) from the three Nagongera mothers sequenced at step d. were raised to adulthood. Genomic DNA was extracted and sequenced from individual males g. and h. Female F1 from each of the six mothers were raised to adulthood. RNA was extracted and sequenced from a pool of ten F1 females three to five days after eclosion.

**Table 1:** Crosses between Kisumu and Nagongera and RNAseq summary statistics.

| RNAseq<br>sample name | Cross<br>name | Mother<br>origin | Mother-Sanger code | Father<br>origin | Father –<br>Sanger code | Total reads<br>counted<br>(F1 RNAseq) | Percent of reads<br>mapping to<br>*Anopheles*<br>*gambiae* PEST<br>genome |
| --- | --- | --- | --- | --- | --- | --- | --- |
| Wilding_1 | B1 | Nagongera | failed QC | Kisumu | unknown | 18,496,263 | 88.6% |
| Wilding_2 | B3 | Nagongera | failed QC | Kisumu | unknown | 38,526,717 | 87.1% |
| Wilding_3 | B5 | Nagongera | AC0382-C | Kisumu | AC0416-C | 28,969,694 | 87.7% |
| Wilding_4 | K2 | Kisumu | AC0300-C | Nagongera | AC0406-C | 6,400,460 | 85.5% |
| Wilding_5 | K4 | Kisumu | AC0317-C | Nagongera | AC0398-C | 46,229,505 | 86.3% |
| Wilding_6 | K6 | Kisumu | AC0334-C | Nagongera | AC0398-C | 29,491,773 | 86.9% |

### Sequencing

RNA was extracted from pools of 10 female F1 progeny from each of the six crosses 3-5 days after eclosion using RNAqueous4PCR total RNA isolation kit (Invitrogen). RNA quality and quantity was checked to be adequate for library preparation using the Agilent Bioanalyser profile and Qubit® 2.0 Fluorometer. Total RNA libraries were prepared for Illumina paired-end indexed sequencing according to the Illumina TruSeq RNA sample preparation v2 guide [37]. cDNA libraries were bar coded, pooled and sequenced using the Illumina HiSeq1500 platform, with 100bp paired end reads.

DNA was extracted using the Qiagen DNeasy Kit from the 6 individual mothers that laid viable eggs, all individual potential fathers that were alive following the cross, and individual male F1 siblings (Figure 2) and sequenced using the Illumina HiSeq 2000 as described previously [6].

**Figure 2:**
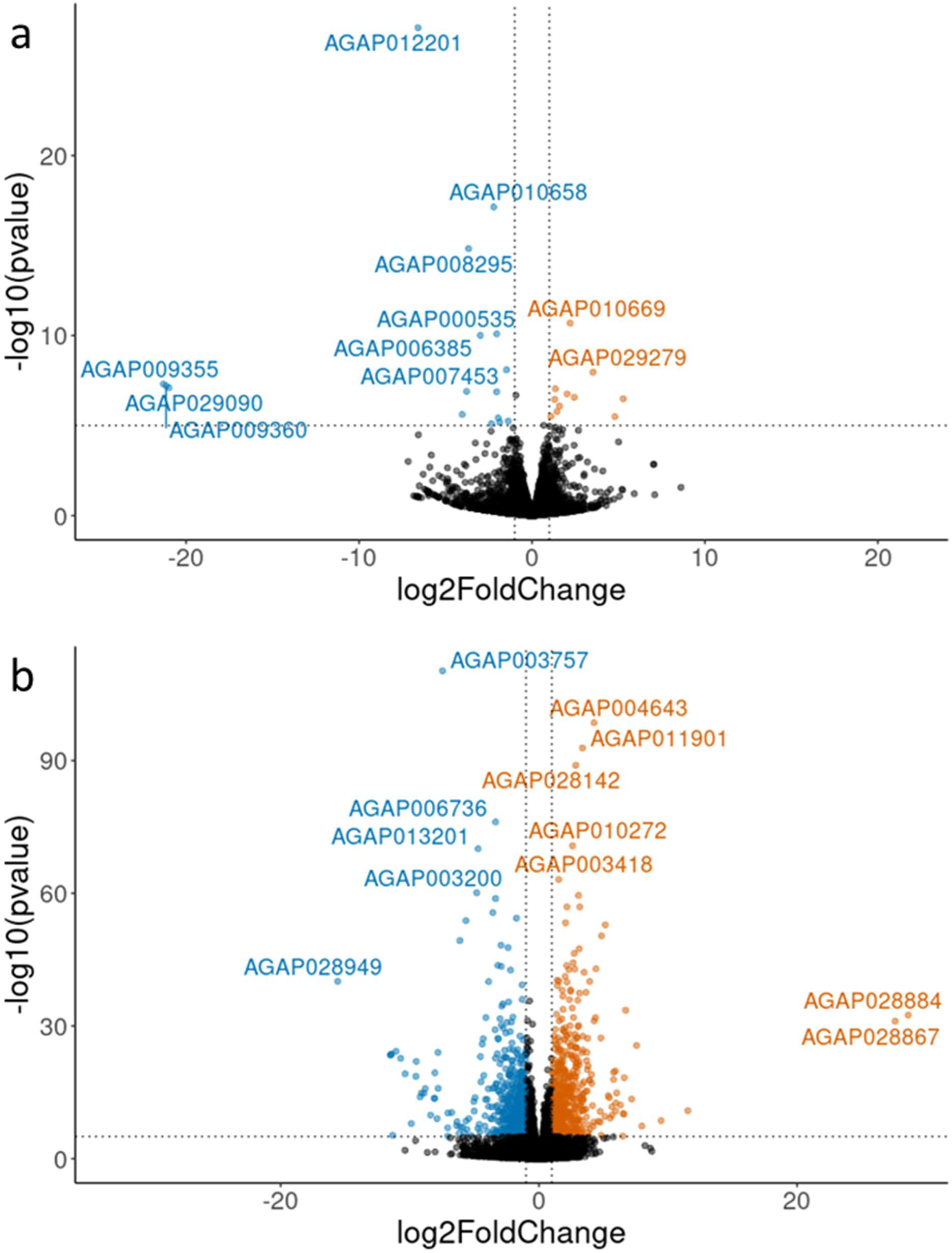
Reciprocal crosses show similar overall gene expression. a. Volcano plot of log2 fold change against -log10 P value comparing gene expression in the F1 progeny of reciprocal crosses between Kisumu and Nagongera strains. Blue points indicate genes downregulated in progeny of Kisumu mothers compared to progeny of Nagongera mothers, orange points indicate genes upregulated in progeny of Kisumu mothers compared to progeny of Nagongera mothers. b. Volcano plot of log2 fold change against -log10 P value comparing gene expression between F1 progeny of Nagongera and Kisumu with the Kisumu parental strain. Blue points indicate genes downregulated in the Kisumu compared to cross progeny, orange points indicate genes upregulated in Kisumu compared to cross progeny.

### Genome analysis

Quality control of genomic sequences for mothers, potential fathers and F1 progeny, and matching of fathers to the progeny using Mendelian Error analysis was performed as described previously [6]. Briefly, for the autosomes for each cross where the mother’s genome sequence was of sufficient quality (K2, K4, K6 and B5), the per progeny sample Mendelian error was calculated for each possible parental pair of the known mother with all potential fathers using scikit-allel [38]. The father in the parental pair that had the smallest median progeny sample Mendelian error on at least 3 autosomal arms was deemed to be the true father.

Scikit-allel [38] was used to analyze the genome of potential fathers including generating a consensus sequence of homozygous sites for Kisumu and Nagongera colonies, and calculating the number of single nucleotide polymorphisms (SNPs) differing between the colonies.

### RNAseq analysis

Samples were compared using the RNA-Seq-Pop snakemake workflow [39] which includes quality control of RNAseq reads using with FastQC [40] and MultiQC [41], read alignment to the Anopheles gambiae PEST reference genome (AgamP4, INSDC Assembly GCA_000005575.1, Feb 2006) using HISAT2 [42], principal components analysis to group samples by gene expression, differential expression analysis using Kallisto [43] with DESeq2 [44], variant calling using Freebayes version 1.3.2 [45] with ploidy 20 (as 10 diploid individuals were pooled in each sample), calculation of population diversity statistics with scikit-allel [38], and gene set enrichment analysis using GSEA [46]. RNAseq data for the Kisumu parental colony, Busia G28 deltamethrin selected colony [39] and Tiefora pyrethroid resistant colony [47] was compared with the cross progeny RNAseq data in this study.

### Selection of SNPs to detect ASE

Detection of ASE relies on sufficient SNP differing between the parents [48]. The number of SNPs differing between maternal and paternal genomes varied (Table 2). For crosses B1 and B3, with parent genomes unavailable, we instead attempted to use parental colony consensus genome sequences for mapping. However, the number of SNPs differing in Nagongera and Kisumu consensus sequences (Table 3) was insufficient for ASE analysis. The low number of differing SNPs may be due to barriers to recombination imposed by the genome structure of *Anopheles gambiae*, which limit the extent to which colonies become inbred, with heterozygosity persisting even after many generations of inbreeding [49]. This contrasts with *Drosophila* where colony genotypes have been used effectively for inference of ASE [50, 51]. Since the mosquitoes were pooled prior to RNA extraction, inferring SNPs from the RNAseq data to detect ASE is problematic, since most SNPs are heterozygous in one or both parents, resulting in a mixture of homozygous and heterozygous SNPs in the progeny, and we would not know at each SNP whether to test for deviation from 1:1 expression or 3:1 expression. To increase the number of SNPs to detect ASE in these pooled samples we therefore used the siblings of the RNA sequenced F1 to infer which SNPs were homozygous but different between the parents (opposite homozygous SNPs). For a Mendelian inherited SNP, opposite homozygous SNP in parents should result in all progeny having heterozygous biallelic SNP. If one or both parents were heterozygous (biallelic) then 50% of the progeny are expected to be heterozygous, and the probability of all the progeny being heterozygous in this situation is given by the binomial probability density function where p=0.5, the number of trials (per SNP) is the number of progeny and the number of “successes” is also the number of progeny. The number of progeny and SNPs per chromosome arm where all progeny were heterozygous is given in Supplementary Table 2. Almost all sequenced siblings were male, the heterogametic sex in *An. gambiae*, so it was not possible to infer SNPs on the X chromosome using this method.

**Table 2:** Number of SNPs in exonic regions differing between the parents that could be used to detect ASE.

| Cross | 2L | 2R | 3L | 3R | X | Y / other | total |
| --- | --- | --- | --- | --- | --- | --- | --- |
| K2 | 13849 | 30134 | 19375 | 20877 | 15543 | 0 | 99778 |
| K4 | 33139 | 35048 | 19074 | 22405 | 15250 | 0 | 124916 |
| K6 | 30303 | 30811 | 14534 | 17801 | 15601 | 2 | 109052 |
| B5 | 29627 | 36203 | 20865 | 25414 | 10539 | 1 | 122649 |

2L, 2R, 3L, 3R and X are the chromosome arms. The numbers are the number of SNPs in exonic regions that are homozygous in the parents of each cross but differ between the parents. These sites should be heterozygous in all cross progeny.

**Table 3:** Number of SNPs differing between Kisumu and Nagongera colony consensus sequences.

| Chromosome | SNPs |
| --- | --- |
| 2L | 2575 |
| 2R | 13507 |
| 3L | 6216 |
| 3R | 2949 |
| X | 10552 |
| Total | 35799 |

“SNPs” shows the number of SNPs differing across all sites in the consensus genome sequences of Kisumu and Nagongera colonies

The suitability of these SNPs to detect ASE was first tested by comparing the SNPs inferred from the siblings to those inferred from the parents, where parental genotypes were known (Supplementary table 3). Most SNPs were shared between the parents and siblings, with only a small fraction (<7.5%) unique to the siblings. The total number of usable SNPs from the siblings was lower than those in the parents: a reduction of between 6% and 16%. Per SNP ASE was calculated as counts at reference SNP/counts at reference SNP plus counts at alternative SNP for the sibling SNPs, and as counts at maternal SNP/ counts at maternal plus paternal SNP for parent SNPs. The mean per SNP ASE for SNPs with at least 10 reads counted remained between 0.49 and 0.51 confirming that the SNPs are likely to be heterozygous in the F1 and that mapping bias is not a problem for this dataset. The standard deviation of per SNP ASE increased very slightly (from 0.12-0.14 to 0.13 to 0.15). For crosses B1 and B3, the per SNP ASE was in line with the other crosses with mean 0.5 and standard deviation 0.13 and 0.15 respectively. However, when the per SNP ASE was plotted against the total counts at each SNP, a cluster of SNPs a with high ASE and expression was seen for the sibling but not parent inferred SNPs (Figure 3, Supplementary Figure 3). SNPs in crosses K2, K4 and B5 that were unique to siblings and with high ASE and high expression were therefore removed from the analysis for crosses B1 and B3 before inferring per gene ASE.

**Figure 3:**
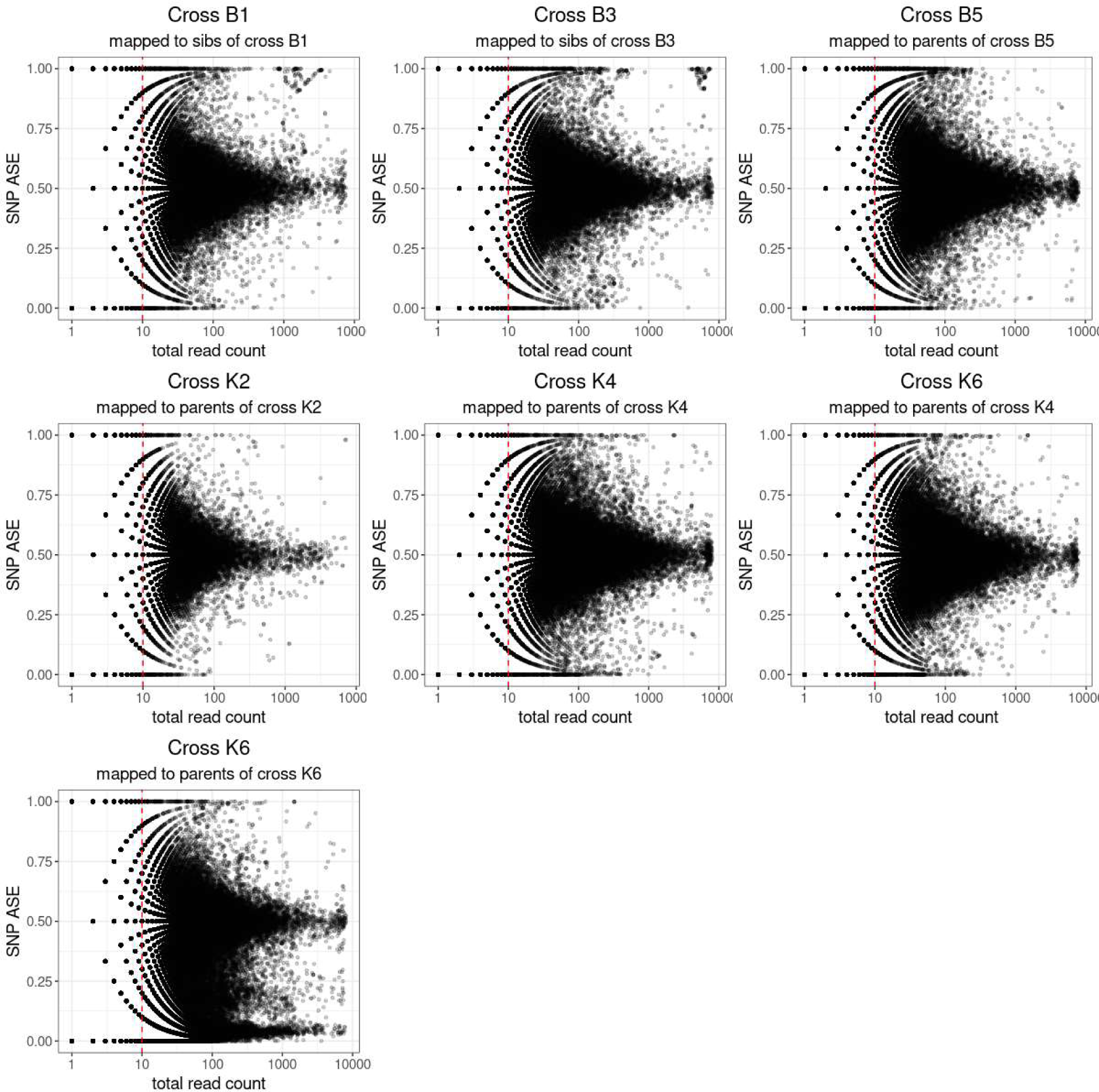
ASE in progeny of crosses between strains. Plot of total read count against ASE at each SNP. Cross and source for the SNPs used to map and count reads are indicated at the top of each plot. Cross K6 is shown with ASE inferred using SNPs from both the parents of cross K4 and for the initially assumed parents of cross K6.

### Analysis of allele specific expression (ASE)

For parent-based mapping, SNPs that were homozygous for different alleles in the parents were used to distinguish whether reads originated from a maternal or paternal allele. Due to the pooling of 10 females for RNA sequencing it was not possible to use parental heterozygous SNPs, since tests of ASE are based on deviation from an expected 1:1 expression ratio and not the 1:3 or 3:1 ratio that would result from one heterozygous and one homozygous parent. For sibling-based mapping, SNPs that distinguished maternal and paternal alleles on the autosomes were inferred from the genome sequences of the male siblings, using the following criteria: SNPs must be heterozygous and biallelic in all siblings where data was available. To eliminate potential low-quality SNPs, if five or more siblings had missing data at a SNP it was not used. Since F1 genome data was unphased, it was not possible to assign SNPs to either the Kisumu or Nagongera parent using this method. Analysis of F1 RNAseq data for ASE was performed using ASEReadCounter* [52], which is based on ASEReadCounter [53], using individual parent genomes where available for each cross to assign reads to parental alleles (options –vcf_mat, -vcf_pat). Where only sibling inferred SNPs were known, option –vcf_joind was used on the unphased inferred F1 VCF file.

The properties of SNPs inferred from parents and siblings were compared by inferring parental pseudogenomes and running ASEReadCounter* with both sets of SNPs. SNP level ASE was calculated for all SNPs with >=10 total counts. SNPs which were inferred from siblings and not from parents, which had >100 counts and ASE < .25 or > .75 were removed from the sibling inferred SNPs for subsequent analysis.

Due to overdispersion, RNAseq data is not binomially distributed but can be better described by a beta binomial distribution. The distribution of counts data and overdispersion parameter rho for the beta binomial distribution was calculated using the counts data for each cross, including the counts for the SNP with the maximum counts in each gene (with minimum 10 counts), using the R package glmmTMB [54], which uses maximum likelihood estimation to fit binomial and beta binomial model to the data for each cross. Models were compared using ANOVA in base R [55].

To obtain a gene level measurement of ASE, the R package MBASED [56] was used to aggregate SNP level counts to gene level ASE by pseudo phasing SNPs in each gene such that the SNPs with the higher read count at each variable site are combined into the major allele, with frequency between 0.5 and 1. Only sites with at least 10 counts were used. For SNPs inferred from the siblings of the RNAseq pool, MBASED was run assuming SNPs were not phased. Where phase was known from the parents, MBASED was run in both phased and non-phased mode to measure the impact on power to detect loci with ASE. In all cases the same seed (988482) was set prior to running 10^6^ simulations in which the total read count at each SNP position in each gene are kept constant, but reference allele counts are drawn from a null distribution beta binomial (mean 0.5 x total count, rho 0.038). The P value is the proportion of simulations in which the major allele frequency is greater than or equal to the observed major allele frequency. For genes with multiple SNPs available to infer ASE, the P value for heterogeneity (Phet) between the level of ASE at each SNP within a gene was additionally calculated. Low P het indicates possible isoform specific ASE. False discovery correction was applied to both P values for ASE and Phet with a nominal rate of 5% [57]. For SNPs inferred from parents, allelic count tables were additionally analyzed using Qleelic [52] with a binomial test of the null hypothesis of equal numbers of transcripts arising from maternal and paternal alleles. ASE was considered significant at p=0.05 following Bonferroni correction. Genes with fewer than 10 read counts were excluded. As technical replicates were unavailable, an exploratory solution for correcting overdispersion was splitting reads into groups (pseudoreplicates K4 14 groups, K2 three groups, K6 nine groups, B5 nine groups). This produced a consistent quality correction constant (QCC) value of 1.56 for QCC correction. R packages UpSetR [58], stats [55] and ggplot2 [59] were used for regressions and plots.

### Enrichment analysis

Hypergeometric tests for enrichment of gene ontology terms, Pfam protein domain enrichment, and KEGG pathways were implemented using python [60]. The test set for each cross was the set of genes showing significant ASE following FDR correction, and the “universe set” was the set of genes for each cross that had at least 10 reads containing SNPs distinguishing maternal and paternal alleles (for these genes ASE would have been detectable if present).

The evolutionary age of genes was taken from the Princeton Protein Orthology Database PPODv4_PTHR7-OrthoMCL [61], with age estimates based on Wagner parsimony. A gene age enrichment test was implemented using the age_enrichment.py script in ProteinHistorian [62], using option ‘-a ignore’ to ignore the proteins not present in the database. P values were obtained using a Mann Whitney U test for difference in overall age distribution between sets and Fisher’s exact test for difference in fraction of proteins in each age in the set.

The null hypothesis, the proportion of genes showing significant ASE does not differ between autosome arm, was tested using a two-sided Fisher’s exact test (for those crosses where the counts were too high for efficient computation P value was simulated using 2000 replicates).

To assess presence of CNV we used the CNV calls for Ag1000g phase 3, which applied a Gaussian hidden Markov Model (HMM) to the normalized windowed coverage data for each genotyped individual using the methods described in Lucas et al 2019 [21]. Copy number variation was deemed possible if any of the parents and F1 siblings from crosses B5, K2, K4 or K6 (total 63 individuals) had a copy number call other than 2 on the autosomes. As ASE could not be inferred on the X chromosome for crosses B1 and B3 we excluded it from the analysis. The father of cross B5, AC0416-C had high variance in copy number calls and was therefore removed from CNV analysis.

To assess selective sweeps, lists of genes either showing ASE or having SNPs to detect ASE but not showing ASE were compared to a database of regions under selection in the *Anopheles* selection atlas [63] as defined by H_12_ signal [64] both across the range of *Anopheles gambiae* and restricted to Uganda using a custom Python script (https://github.com/azurillandfriend/sweeps_ASE.git).

### Prediction of *cis*-regulatory modules

*Anopheles cis*-regulatory modules (CRMs) were predicted using SCRMshaw [65–68]. Briefly, *Drosophila* CRMs which drive gene expression in the tissue of interest were downloaded from the Redfly database (v9.6.0, database updated 02/01/2023) [69, 70],with max size 2000. Only non-overlapping sequences (>100bp) were included. Syntenous regions from related *Drosophila* species (putatively containing the equivalent CRM) were added to the training data set using liftOver at https://genome.ucsc.edu/cgi-bin/hgLiftOver [71] as described in [66]. SCRMshaw was trained using this augmented training set and a 10x bigger set of non-CRM non-exonic regions. Repeats in CRMs, non-CRMs and the target *Anopheles gambiae* PEST genome were masked using repeat masker [72]. Existing training sets for adult peripheral nervous system, embryonic and larval excretory, and embryonic/ larval Malpighian tubules were downloaded from GitHub (https://github.com/HalfonLab/dmel_training_sets). These were compared with the 2272 Tn5 transposase hypersensitive sites identified by Ruiz et al [73].

## Results and Discussion

### Crosses

Three females from each reciprocal cross between the Kisumu and Nagongera strains laid viable eggs (Figure 1, Table 1). For four crosses the father was identified by minimizing median Mendelian error (Supplementary Table 1, but for the other two crosses the mother failed sequencing quality control and none of the sequenced putative fathers were a good match. We presume the true fathers for these crosses died during the experiment, precluding extraction of good quality DNA.

RNAseq statistics for the pooled F1 females from each of these six crosses are shown in Table 1. The number of reads recommended for 60% power to detect ASE at 1.5 fold is 500 per gene [48], which for the 13,796 annotated genes in *Anopheles gambiae* would require around 6.9x10^6^ reads. Five crosses exceeded this value with one cross, K2, having slightly fewer (6.4 x10^6^reads). The proportion of reads mapping to the *Anopheles gambiae* genome was similar for all crosses (mean 87.0%, standard deviation 1.1%).

### Between pools gene expression comparison

Comparison between reciprocal crosses revealed similar gene expression in the F1 progeny, with just 38 genes significantly downregulated and 38 genes significantly upregulated in the F1 between reciprocal crosses at P_adj_ ≤ 0.05 (Figure 2a). When comparing only F1 crosses in PCA, B1 seemed to be an outlier in terms of expression (Supplementary figure 1a). However, when compared with other colonies of *Anopheles gambiae* s.l., the F1 progeny formed a distinct group (Supplementary figure 1b).

F1 gene expression differed from the Kisumu (parental) colony, with a total of 1362 genes upregulated and 1265 genes downregulated in the F1 compared to the Kisumu colony (Padj ≤ 0.001) (Figure 2b). RNAseq data was unavailable for the Nagongera colony, which no longer exists. Gene set enrichment analysis of the differentially expressed genes indicated that genes with Gene Ontology (GO) terms associated with odorant binding, olfactory receptor activity, sensory perception of smell, response to stimulus, detection of chemical stimulus involved in sensory perception of smell, structural constituent of cuticle and signal transduction were downregulated in the F1 compared to Kisumu, whereas those with GO terms associated with translation, ribosome, mitochondrion, structural component of ribosome and serine type endopeptidase activity were upregulated in the F1 compared to Kisumu (all at P adjusted 0.05). KEGG pathway analysis showed significant upregulation of ribosome, citrate cycle (TCA cycle) and oxidative phosphorylation in the F1 compared to Kisumu, and no KEGG pathway significantly downregulated.

### ASE inference

For the four crosses where the parent was known, per gene ASE was calculated as reads mapping to maternal genome/total reads mapped at that gene. In addition the major allele frequency at each gene was calculated for all crosses. Read count data were fitted to binomial and beta binomial models. ANOVA indicated that the beta binomial model was a better fit to the data (P < 2.2e-16). The dispersion parameter phi(disp) for the beta binomial distribution of counts data for crosses B1, B3, B5, K2, and K4 was estimated at 25.8, giving a rho 1/(1+phi(disp)) of 0.038. Per gene ASE statistics for all crosses are shown in Table 4. Due to variations in parental genome and sequencing depth the power to detect ASE varies between crosses, so it is not possible to infer whether the proportion of genes with ASE is different between the crosses. Three crosses showed similar numbers of genes with maternal or paternal ASE except cross K6 which showed extreme paternal ASE. This could either be due to K6 showing a very different pattern of ASE to the other crosses, or a sampling error occurred so that the RNA sequenced samples were not from the same cross as the parents and siblings. We therefore checked the effect of using SNPs from the wrong parents to infer ASE. Using SNPs from non-matching parents resulted in inflated ASE estimates (Supplementary Table 4). Using SNPs from the parents of cross K4 to infer allelic counts for cross K6 shifted mean imbalance back to 0.5 (figure 3, Supplementary Table 4). Furthermore, when SNPs called from the RNAseq data were compared, the progeny of cross K4 and K6 were extremely genetically similar and clustered closely in PCA for all chromosomal arms (Supplementary Figure 4), suggesting they may have hatched from two egg batches from the same cross. Gene level ASE for cross K6 was therefore inferred using SNPs from K4 parents.

**Table 4:** Number of genes showing ASE in each cross.

| Cross | Genes showing ASE | Total genes with SNP(s) and sufficient counts to detect ASE |
| --- | --- | --- |
| B1 | 327 | 6486 |
| B3 | 324 | 6065 |
| B5 | 823 | 7537 |
| K2 | 255 | 5185 |
| K4 | 979 | 8166 |
| K6 | 757 | 7852 |
| All (intersection) | 13 | 2934 |

ASE detected with FDR of 5%
The totals for B1 and B3 are after the removal of SNPs which were inferred from siblings and not from parents, which had >100 counts and ASE < .25 or > .75

Genes showing ASE showed some overlap between the different crosses (Figure 4). At the most conservative estimate, 13 genes showed ASE in all crosses. This exceeds the overlap expected by random genes showing ASE, as in 100,000 simulations of randomly drawing the observed number of significant genes from the set of genes where SNPs were available to detect ASE in all crosses, the maximum overlap was 1. Most genes showing ASE were unique to individual crosses (Figure 4), but there were many genes showing ASE in combinations of multiple crosses that may also be under consistent differential *cis*-regulation between Tororo and Nagongera colonies, Table 5 shows the 115 genes with significant ASE in at least 4 out of the 6 crosses. Directionality of ASE (maternal or paternal bias) was inferred for crosses B5, K2, K4 and K6 (Supplementary Table 7).

**Figure 4:**
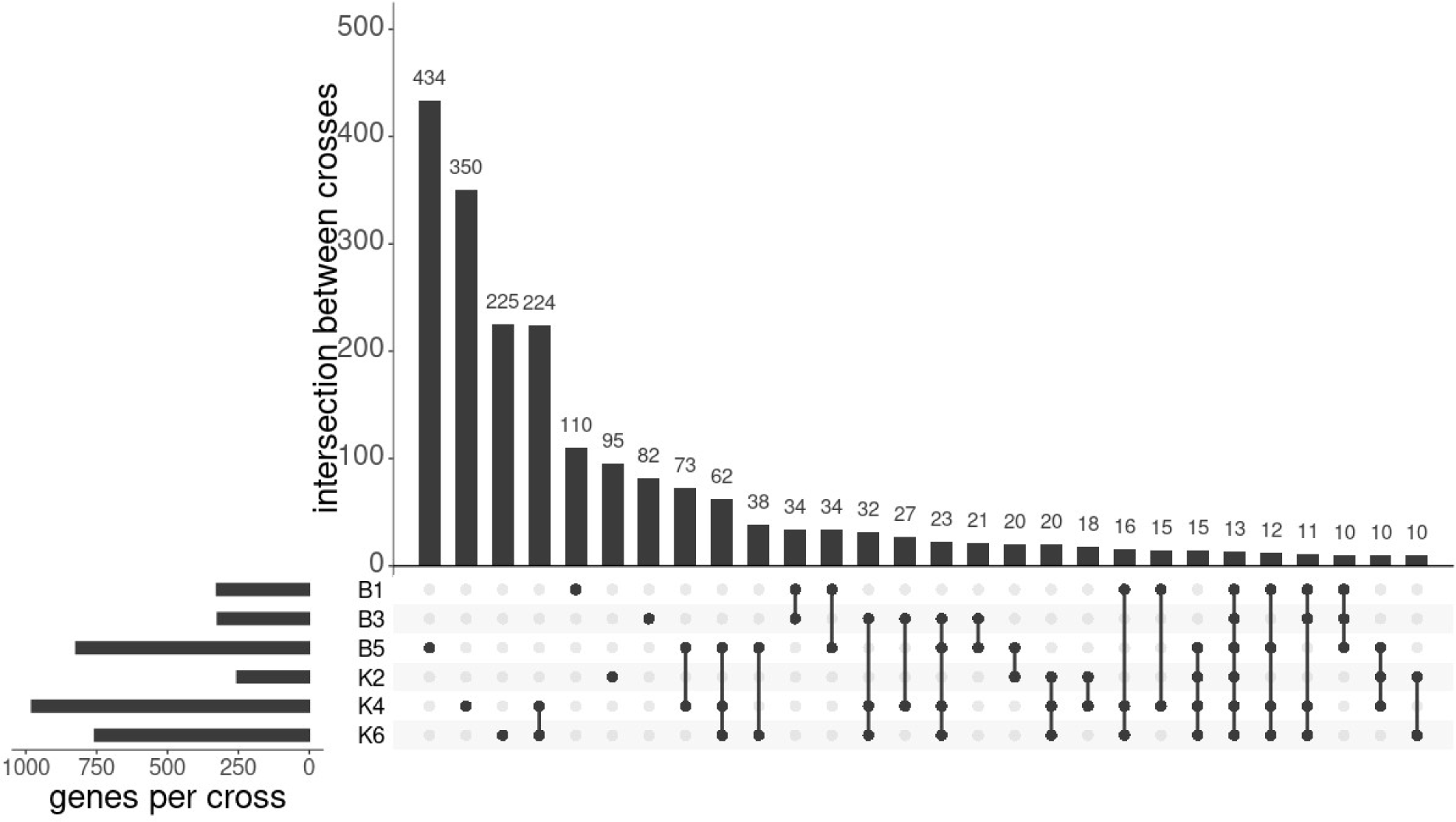
Intersection of genes showing ASE between crosses. UpSet plot with the number of genes showing ASE for each cross and the intersection of these genes between crosses. Only the first 28 sets of overlaps are shown.

**Table 5:**
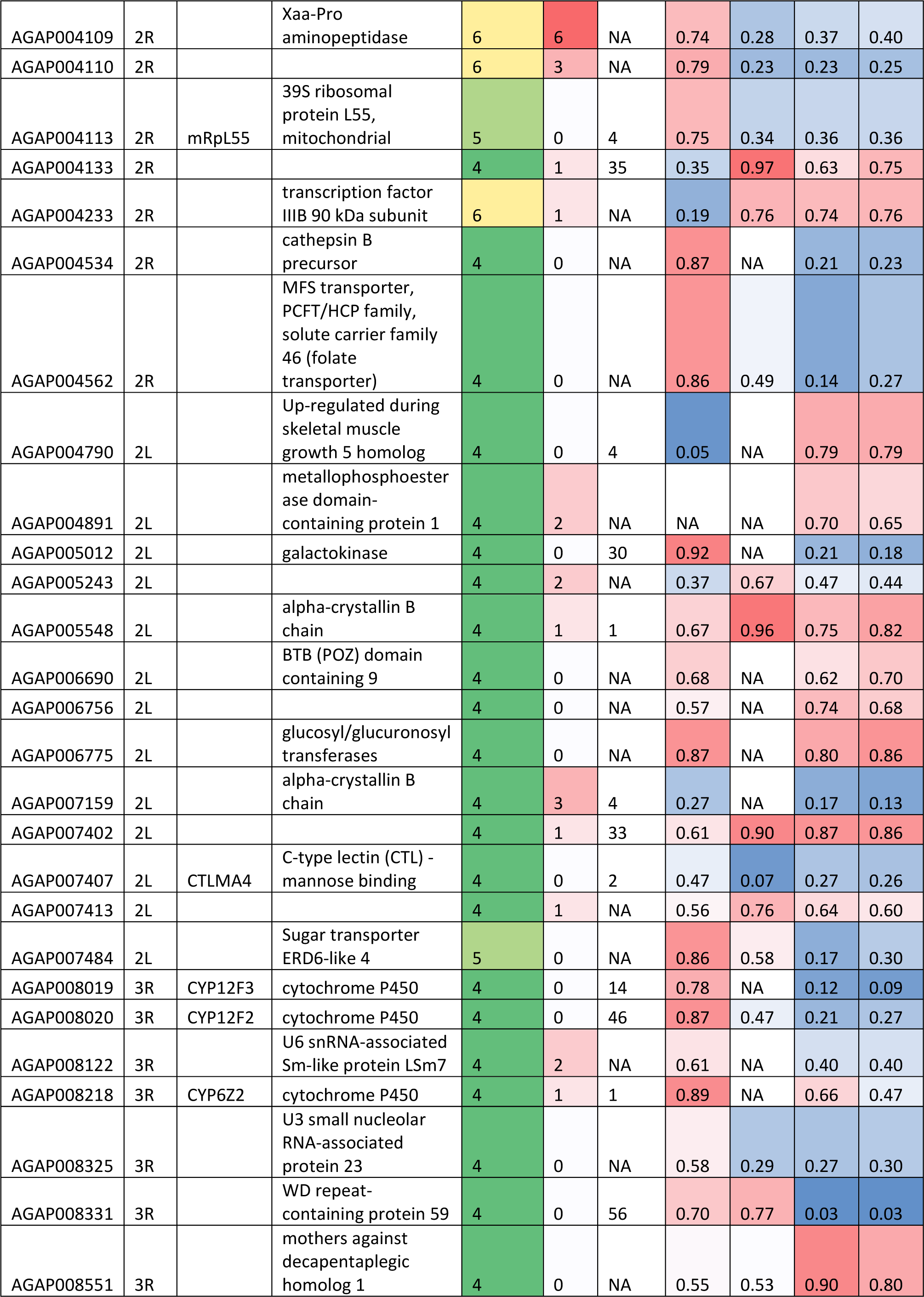
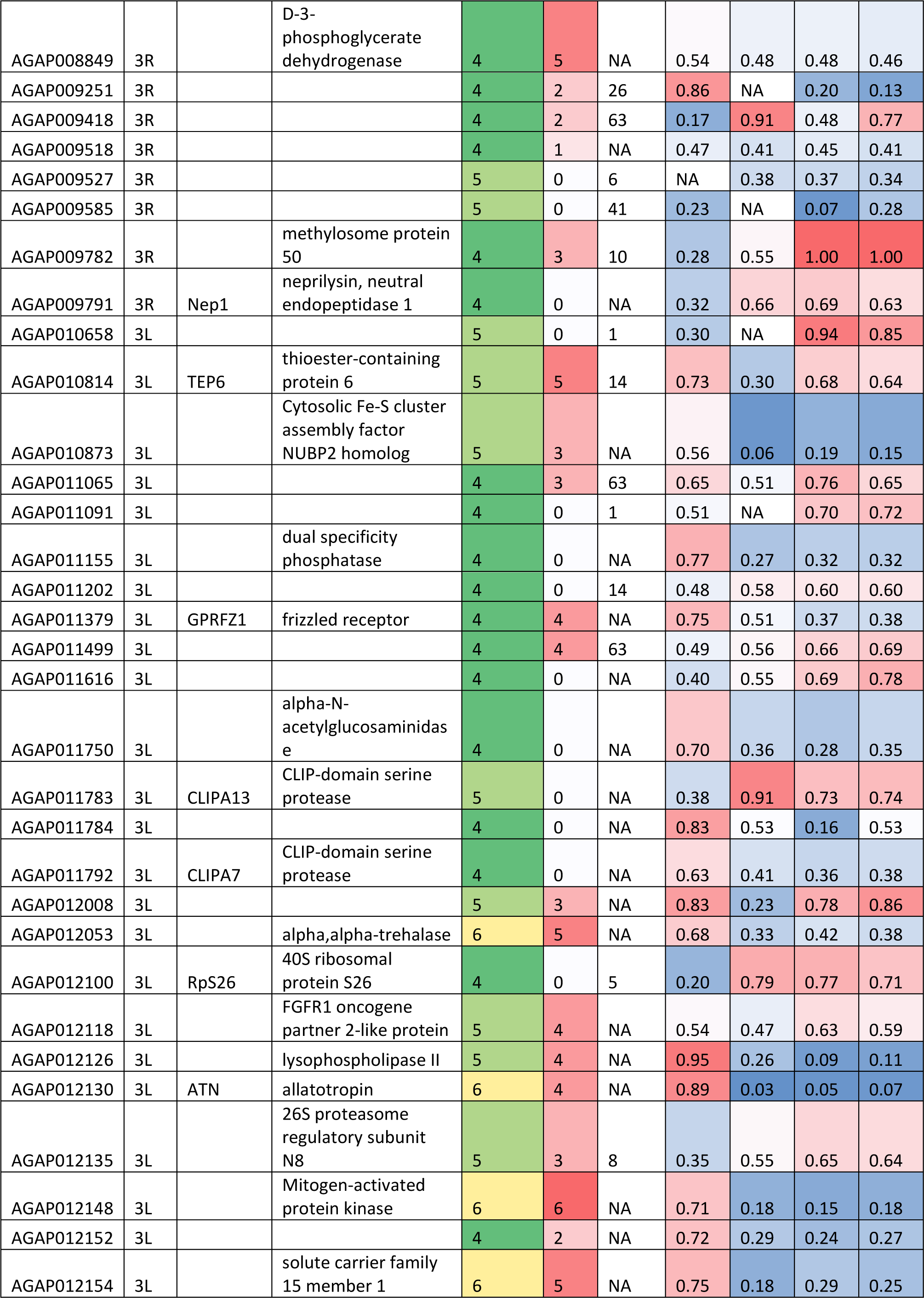
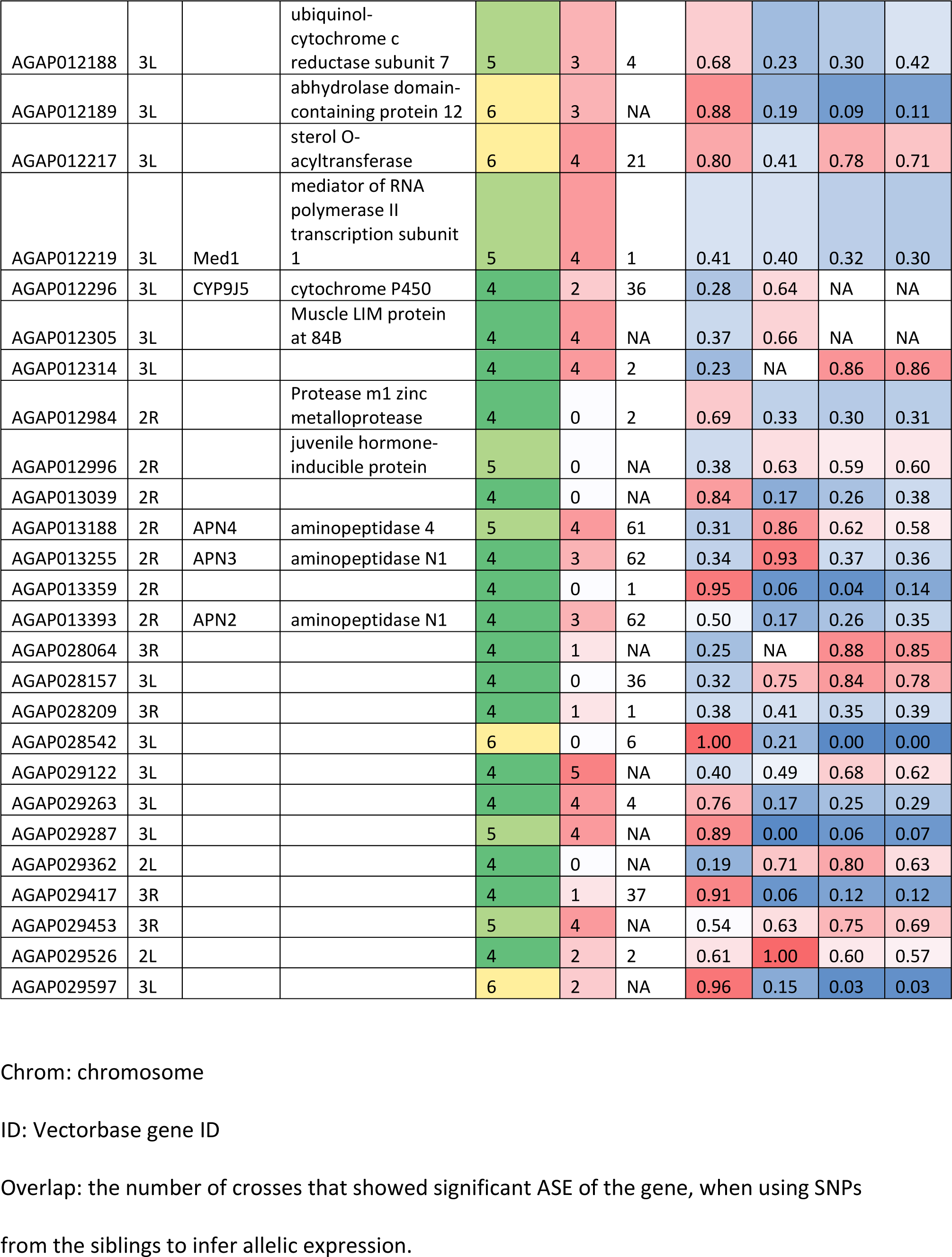
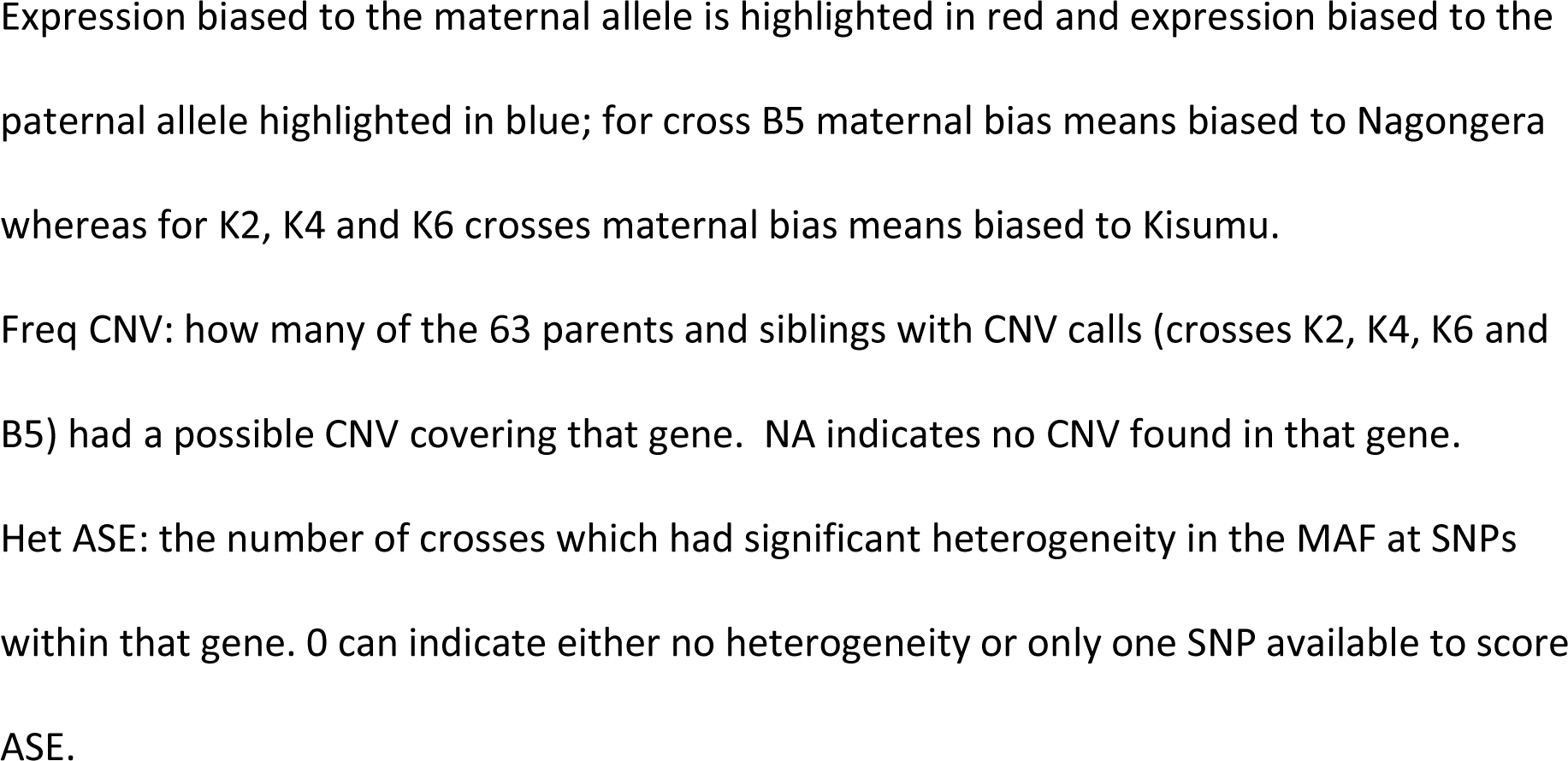
Genes showing ASE in at least four crosses.

Analysis of gene ages that showed or did not show ASE revealed an enrichment of younger, *Anopheles* specific genes showing significant ASE (Figure 5 and Supplementary Table 5). The same trend was seen in all crosses.

**Figure 5:**
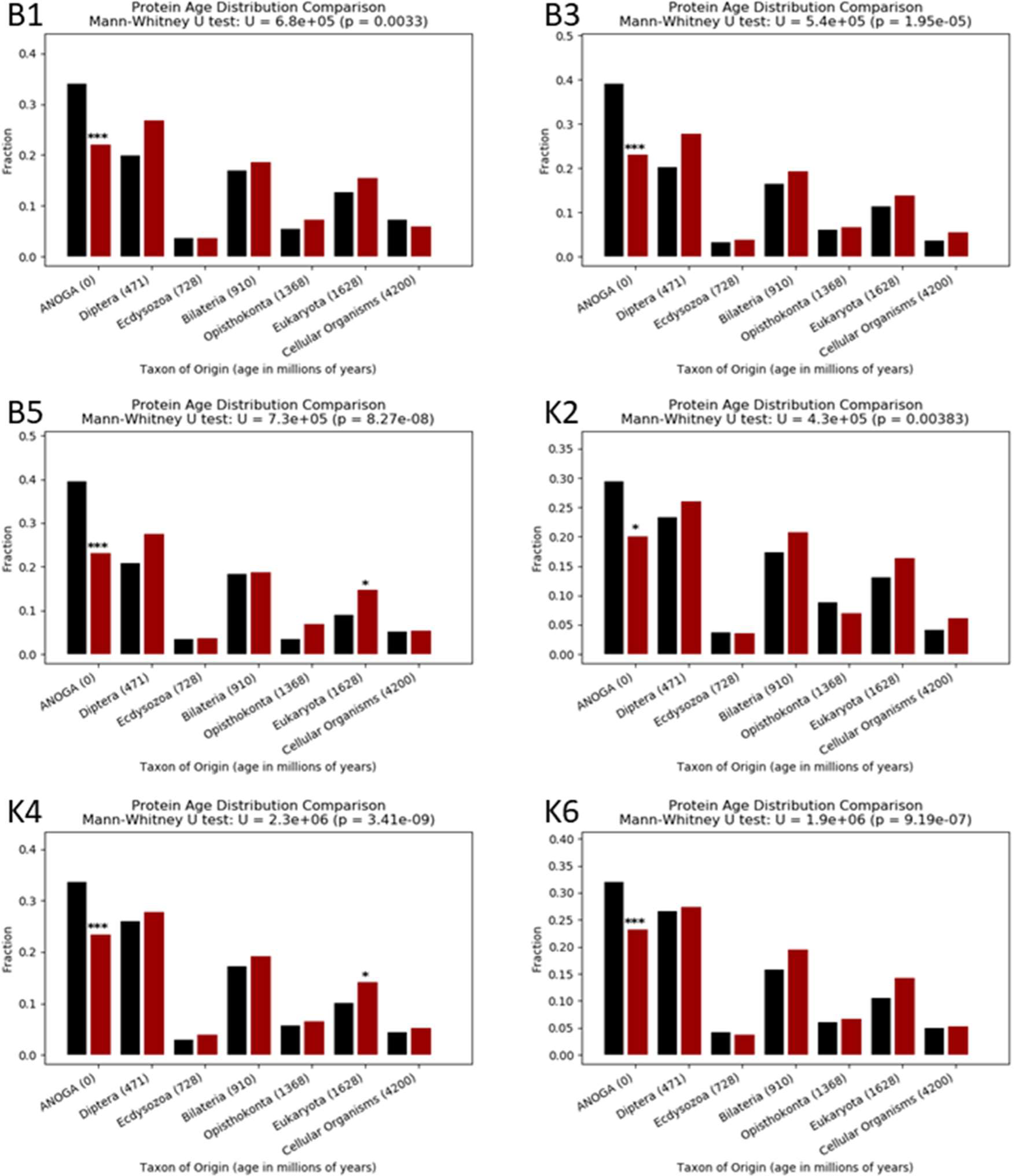
Ages of genes showing ASE or not showing ASE. bar plots comparing the age of genes which showed ASE or did not in the progeny of six crosses between Nagongera and Kisumu strains, using the Wagner parsimony method. The cross name is indicated at the top left of each plot, Fisher’s exact test P values for the difference in fraction of genes in each age between genes showing ASE (red bars) and genes not showing ASE (black bars) are displayed above the bars; *: 0.001<P<0.05, ***:P≤0.001.

Genes showing ASE showed an unequal distribution along chromosomal arms compared to genes not showing ASE (Supplementary table 6, Figure 6, Supplementary figure 2). This trend was significant for all crosses except for K2. The trend was for slight enrichment on 3L. For the 4 crosses where parents were available, this was also compared with the X chromosome, which had a lower proportion of ASE/total detectable genes (Figure 6, Supplementary figure 2). ANOVA indicated both chromosome and cross explained the variance in the proportion of ASE/total detectable genes per chromosome arm (P < 0.05).

**Figure 6:**
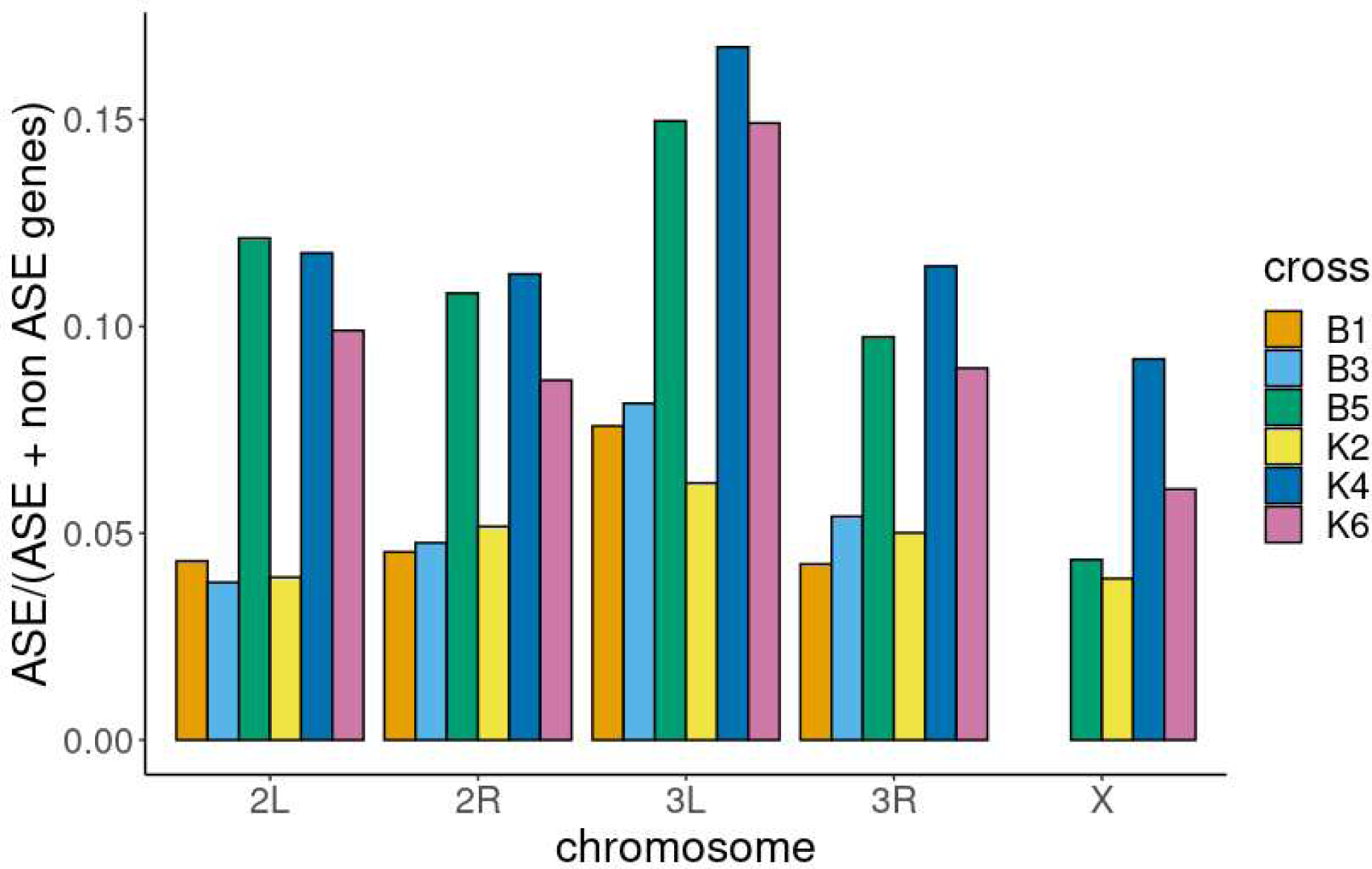
proportion of genes showing ASE versus genes with sufficient SNPs to detect ASE along each chromosome arm. Bar plot of the proportion of genes showing allele specific expression/ total genes containing suitable SNPs to detect ASE per chromosome arm. Colours indicate the different crosses. X chromosomal SNPs were not available for crosses B1 and B3

### Copy number variation

True ASE is caused by *cis*-regulation, but copy number variation of the expressed gene could lead to apparent ASE if there are different numbers of gene copies containing the SNPs used to count reads; e.g. if two copies of a duplicated gene with total of three copies bear one SNP and the third the other SNP, apparent ASE would be inferred without differential *cis*-regulation. We therefore checked for CNV in the parental and F1 sibling genomes at the genes that showed ASE in at least four out of six crosses, and in the genes which contained sufficient SNPs to infer ASE but expression appeared to be in balance. A total of 60/114 (53%) autosomal genes showing ASE contained possible CNV. For genes that did not show ASE and had available SNPs to infer ASE in all crosses, 481/ 1333 (36%) had possible CNV. A two-sided Fisher’s exact test rejected the hypothesis that the odds ratio is 1 (odds ratio 2.0, P=0.0006). This suggests that for some genes CNV may be contributing to the observed ASE. The number of individuals with a possible CNV in each gene varied between the genes (Table 5).

### Overlaps between ASE genes and other datasets

The list of 115 genes showing ASE in at least four crosses was compared with published gene expression data. All other available *Anopheles* expression data is based on between sample comparison and represents a mixture of *cis* and *trans* regulation, whereas ASE data is specific to *cis* regulation and excludes any *trans* regulation. Therefore even for extremely similar ASE and between sample comparison datasets only *cis*-regulated genes are expected to overlap.

Genes showing ASE in at least four crosses were compared with genes with consistently high median fold change in a meta-analysis of 35 experiments comparing RNAseq data between *Anopheles gambiae* s.l. and *Anopheles funestus* strains [60], and with genes showing significant fold change with consistent directionality in microarray data comparing susceptible and resistant populations of *Anopheles coluzzi* [74]. For the RNAseq metadata set, the genes with the top 5% of median fold changes between susceptible and resistant populations (429/8599 total genes in the dataset), six genes were also present in the ASE gene set. These were AGAP001251 (*Eupolytin*), AGAP008218 (*Cyp6Z2*), AGAP012296 (*Cyp9J5*), AGAP011068 (A*ldose reductase*), AGAP003583 (*L-iditol 2-dehydrogenase*) and AGAP008331 (*WD repeat-containing protein 59*). There was no overlap between ASE and microarray data.

Gene set enrichment analysis did not reveal any consistent patterns across crosses for enrichment of genes showing significant ASE in any GO term or Kegg pathway. Trypsin and Ubiquitin pfam domains were significantly enriched in 4/6 crosses.

Of the gene families previously implicated in metabolic resistance, we observed four P450s with ASE in four of the six crosses: *Cyp12F3* (AGAP008019), *Cyp12F2* (AGAP008020), *Cyp6Z2* (AGAP008212) and *Cyp9J5* (AGAP012296).

The D7 protein family has previously been implicated in bendiocarb resistance in Uganda [25]. AGAP008282 (*D7r2*) showed significant ASE in two crosses (detectable in 4) and *D7r4* in none (detectable in 3). Other *D7r* genes, *D7r1* (AGAP008284), *D7r5* (AGAP008280) also showed significant ASE in two crosses, *D7L1*, *D7L2* and *D7r3* in one cross.

We finally asked whether genes showing ASE were more likely to be in a genomic region that has undergone a recent selective sweep. Out of 115 genes showing ASE in at least 4/6 crosses, 13 were in a swept region, whereas 103 of the 1333 genes showing no evidence of ASE in any of the 6 crosses were in swept regions. A two sided Fisher’s exact test did not reject the hypothesis that the odds ratio is 1 (odds ratio 1.5, P=0.2), suggesting that genes showing ASE are no more likely to be in a swept region than genes that do not.

### CRM prediction

*Drosophila* CRMs for adult midgut, adult Malpighian tubules, larval midgut and legs were used as training data from SCRMshawHD to predict *Anopheles* CRMs operating in the same tissues, together with previously developed training sets for the adult peripheral nervous system, embryonic and larval excretory system and embryonic/ larval Malpighian tubules (https://github.com/HalfonLab/dmel_training_sets). These tissues were selected based on previous studies examining the tissue specific expression of genes involved in insecticide resistance with roles including detoxification and cuticular resistance [75–80]. Training CRM sets used for the first time in this study are shown in table 6, with full sequences at https://github.com/azurillandfriend/traning_sets_IR.git. Oenocyte (FBbt:00004995), cuticle (FBbt:00004970) and adult epidermis (FBbt:00005401) CRMs could not be used as training data due to insufficient experimentally validated CRMs, highlighting the need for more research to identify CRMs in these tissues. The top scoring 250 predictions for each training set and method were combined, producing a total of 4122 unique CRM predictions. 62 of these predicted CRMs were flanked by a gene showing ASE in at least 4/6 crosses (Table 7). In total, CRMs were predicted for 33 of the 115 genes showing ASE in at least 4/6 crosses. 211 predicted CRMs were flanked by a gene showing consistently high median fold change between resistant and susceptible strains (Supplementary table 8). CRMs were predicted for a total of 141 of the 429 genes in this set. The predicted CRMs were also compared with a published dataset of Tn5 transposase sensitive sites in the adult midgut [73, 81]. 50 of the predicted CRMs overlapped with Tn5 transposase sensitive sites identified as *cis*-regulatory elements by Ruiz et al [73], flanking a total of 48 genes. The vast majority of these CRMs (47) were previously predicted by Kazemian [82] despite the different training sets. The CRMs identified in these predictions provide a starting point for future studies to examine genetic variation in Anopheles populations with different insecticide resistance phenotypes, and for validation using reporter assays.

**Table 6:** training CRM.

| Tissue | Anatomy<br>ontology term | Date of<br>search | Total<br>Number of<br>CRM<br><2000bp in<br>Redfly at<br>time of<br>search | Number of<br>CRM<br>meeting<br>criteria (non<br>overlapping) |
| --- | --- | --- | --- | --- |
| Adult Midgut | FBbt:00003138 | 12/12/22 | 12 | 11 |
| Larval<br>Midgut | FBbt:00005825 | 12/12/22 | 16 | 15 |
| Adult<br>Malpighian<br>Tubules | FBbt:00005725 | 12/12/22 | 10 | 9 |
| Leg | FBbt:00004640 | 12/12/22 | 35 | 28 |
| Antennae | FBbt:00004511 | 24/04/23 | 25 | 20 |
| Oenocyte | FBbt:00004995 | 25/04/23 | 3 | 2 |

**Table 7:**
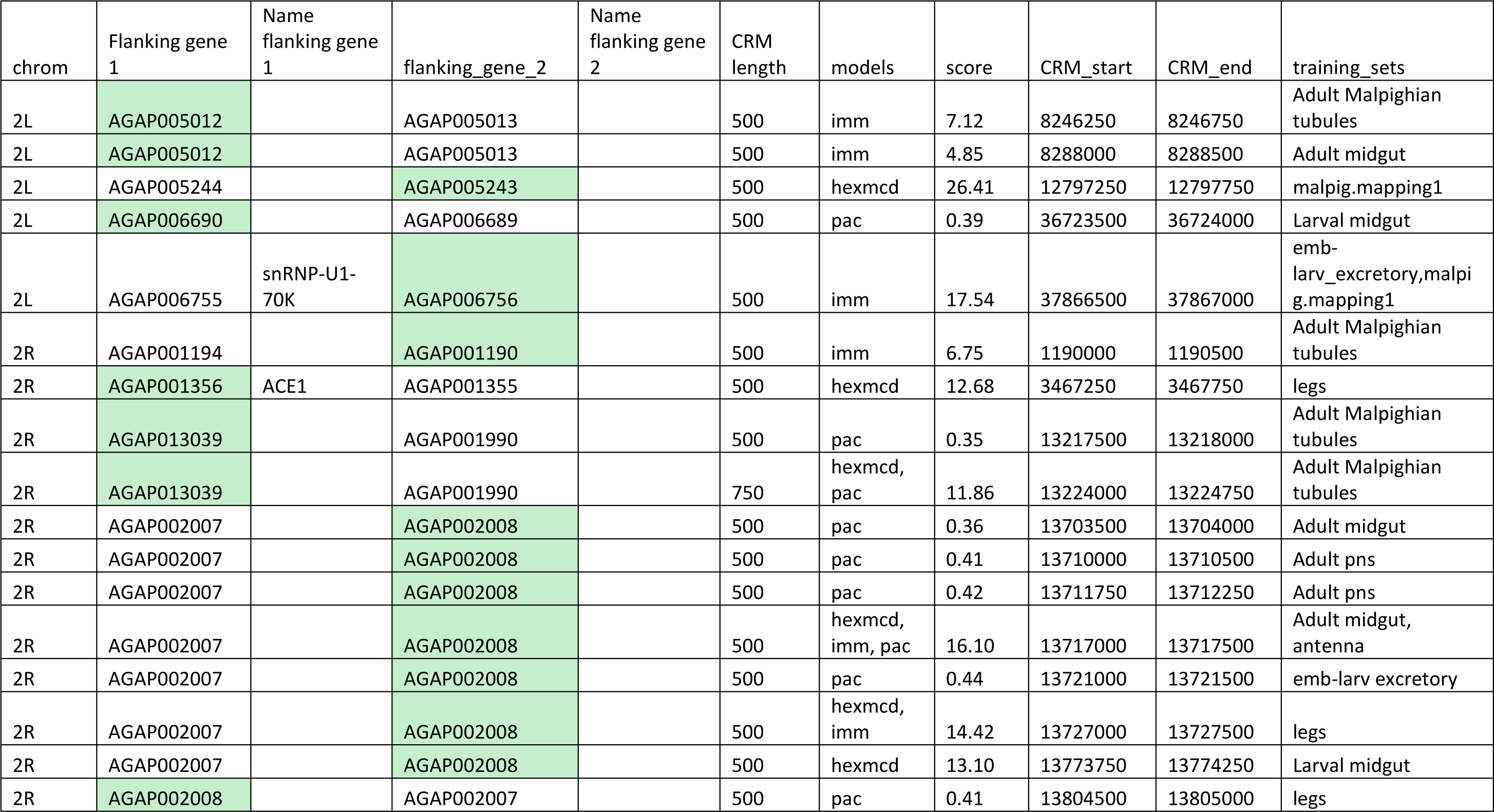
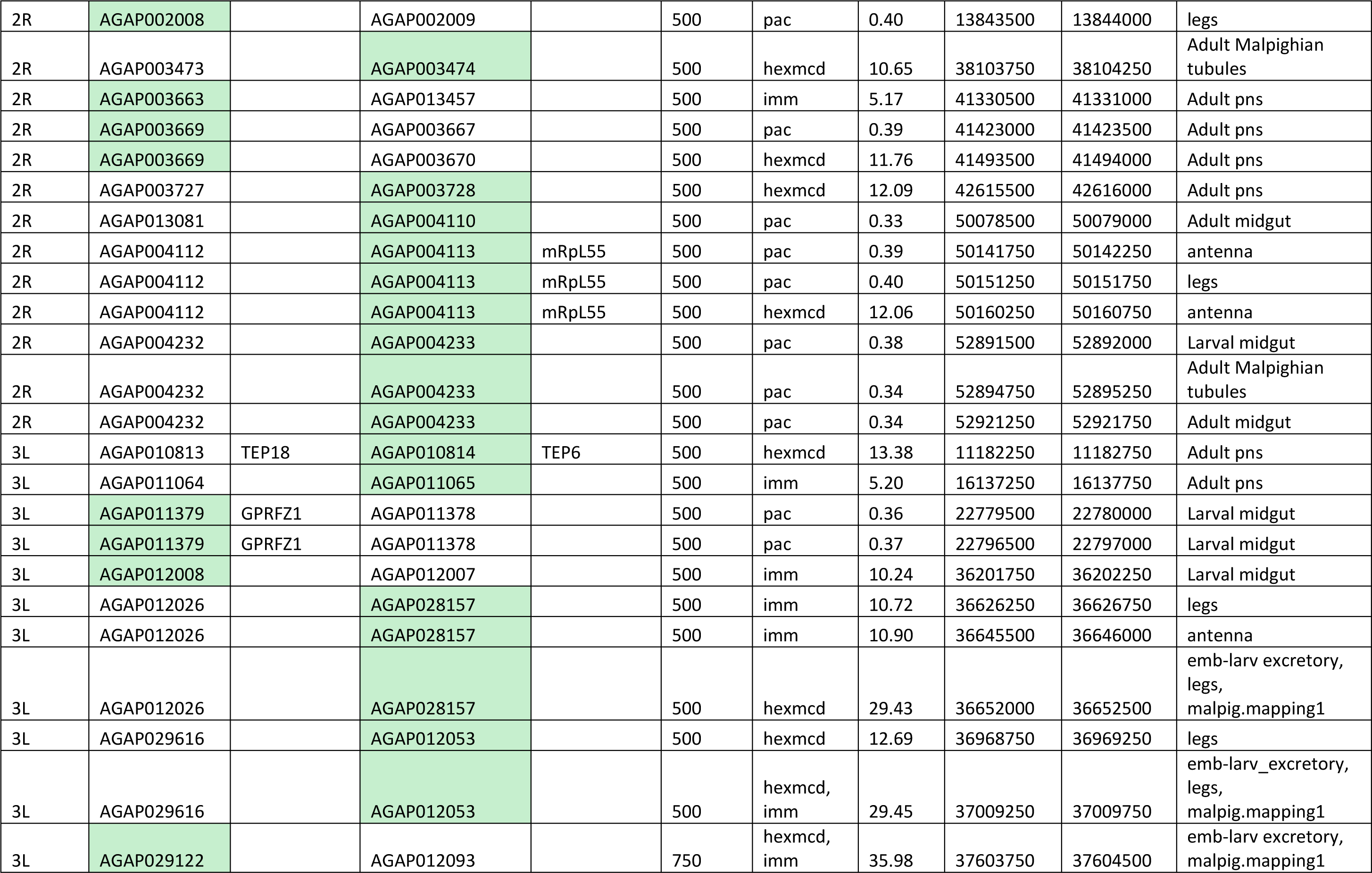
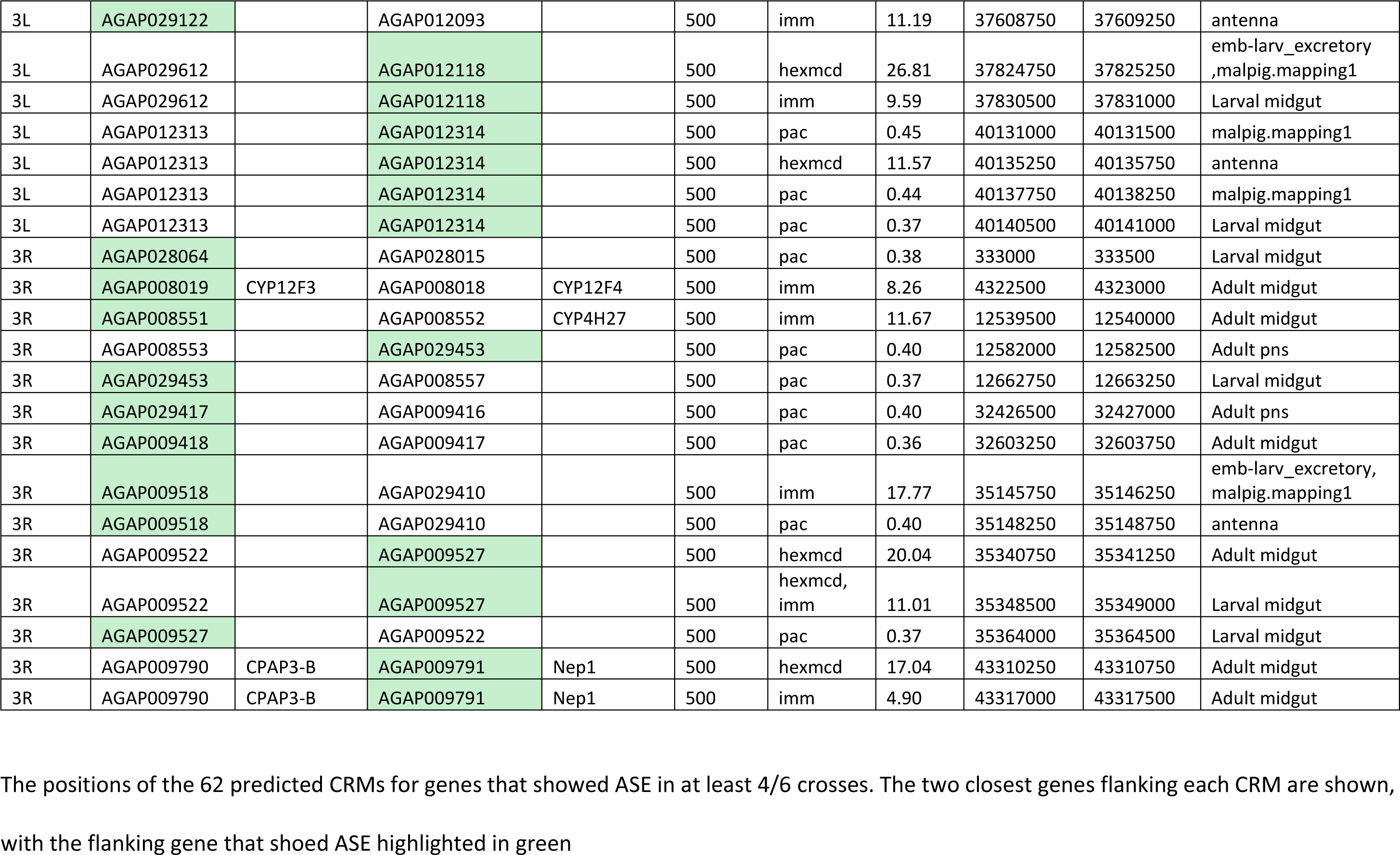
CRM predictions for genes that showed ASE in at least 4/6 crosses.

## Conclusions

This study of allele specific expression in *Anopheles* provides the first evidence that *cis*-regulation of gene expression occurs across the genome and differs between strains in this species.

Sample pooling, while a cost-effective solution for bulk RNAseq of small samples, does limit the power of experiments targeting the study of ASE; in future, to maximize the power to detect ASE, individual samples should be used. RNAseq on whole mosquitoes may have masked tissue specific ASE. The true number of genes showing ASE in individual tissues is likely higher than we observed on the whole mosquitoes analyzed here, since pleiotropic effects may limit the potential for genes to up or down regulated in all tissues simultaneously but permit the evolution of tissue specific regulation [83]. Future studies should target specific tissues of interest. Despite these limitations, we were able to detect genes showing ASE in pooled RNA from whole mosquitoes in crosses between *Anopheles* strains with different carbamate resistance phenotypes, indicating different *cis*-regulation patterns between the strains. Whilst we detected some genes previously implicated in insecticide resistance, there was no consistent enrichment of these or any particular gene ontology term amongst genes showing ASE. This probably indicates that the strains have undergone extensive *cis*-regulatory divergence, affecting both genes involved in insecticide resistance but also genes involved in many other functions. Comparing ASE in progeny of crosses between multiple insecticide resistance and susceptible strains, together with examining gene expression in the parental strains, would enable a more comprehensive survey of the *cis* versus *trans* regulation of insecticide resistance genes. The bias towards younger, *Anopheles* specific genes showing ASE in the F1 suggests there may be a higher degree of *cis*-regulatory divergence between the parental strains for younger genes. It was possible to computationally predict some CRM involved in tissue specific expression, including potential CRM for genes showing ASE and genes previously implicated in insecticide resistance. This was hampered by the lack of good quality training data for the tissues thought to be most relevant to insecticide resistance in adult mosquitoes, highlighting the need for future experimental CRM discovery.

## Data availability statement

1. RNAseq data for all crosses and read counts per SNP are available at Gene Expression Omnibus accession GSE241768.
2. Potential fathers sequencing data are available at European nucleotide archive (accession numbers in Supplementary Table 9).
3. Sequencing data from siblings from crosses B1 and B3 are at European nucleotide (accession numbers in supplementary Table 10)
4. All other nucleotide sequences used are already publicly available in the Phase 3 release of the *Anopheles gambiae* 1000 genomes project, at accession PRJEB42254. For convenience, individual accessions for the parents and F1 male sibs in crosses B5, K2, K4 and K6 are in Supplementary Table 11.

## Acknowledgements

This project was funded by Daphne Jackson Fellowship to N.A.D. sponsored by the Biotechnology and Biosciences Research Council, with RNA sequencing funded by Infravec (www.infravec.eu). Additional support to this work was provided by the National Institute of Allergy and Infectious Diseases (NIAID R01-AI116811) and the Medical Research Council (MR/T001070/1). M.J.D. is supported by a Royal Society Wolfson Fellowship (RSWF\FT\180003).

This study was supported by the MalariaGEN Vector Observatory which is an international collaboration working to build capacity for malaria vector genomic research and surveillance, and involves contributions by the following institutions and teams. Wellcome Sanger Institute: Lee Hart, Kelly Bennett, Anastasia Hernandez-Koutoucheva, Menelaos Ioannidis, Julia Jeans, Paballo Chauke, Victoria Simpson, Eleanor Drury, Osama Mayet, Sónia Gonçalves, Katherine Figueroa, Tom Madison, Kevin Howe, Mara Lawniczak; Liverpool School of Tropical Medicine; Broad Institute of Harvard and MIT: Jessica Way, George Grant; Pan-African Mosquito Control Association: Jane Mwangi, Edward Lukyamuzi, Sonia Barasa, Ibra Lujumba, Elijah Juma. The authors would like to thank the staff of the Wellcome Sanger Genomic Surveillance unit and the Wellcome Sanger Institute Sample Logistics, Sequencing and Informatics facilities for their contributions.

The MalariaGEN Vector Observatory is supported by funding awarded to Dominic Kwiatkowski and Mara Lawniczak from Wellcome (220540/Z/20/A, ’Wellcome Sanger Institute Quinquennial Review 2021-2026’) and funding awarded to Dominic Kwiatkowski from the Bill and Melinda Gates Foundation (INV-001927). The Liverpool School of Tropical Medicine’s participation was supported by the National Institute of Allergy and Infectious Diseases ([NIAID] R01-AI116811), with additional support from the Medical Research Council (MR/P02520X/1). The latter grant is a UK-funded award and is part of the EDCTP2 programme supported by the European Union. The Pan-African Mosquito Control Association’s participation was funded by the Bill and Melinda Gates Foundation (INV-031595).

The authors are grateful for discussions with Louise Cerdeira, Laura Brettell and Shannon Quek, and thank Luciene Salas Jennings and Andrew Carey for providing administrative support to the project.

## Notes

https://www.ncbi.nlm.nih.gov/geo/query/acc.cgi?acc=GSE241768

https://www.omicsdi.org/dataset/omics_ena_project/PRJEB42254

https://www.ebi.ac.uk/ena/browser/view/PRJEB1670

### Competing Interest Statement

The authors have declared no competing interest.

### Summary of Updates

1. The addition of supplementary table 11 which contains ENA accessions for the genome sequence of the parents and F1 male siblings described in the manuscript. 2. The addition of supplementary figure 4 which contains PCA plots. 3. A paragraph of the results and discussion describing a comparison between the results of this study and Isaacs et al 2018 was removed as the methods were too dissimilar for comparison. 4. The paragraph describing the selection of SNPs used to infer ASE was moved to Methods.

