## Supplementary Figures for "Mechanisms of transcriptional regulation in *Anopheles gambiae* revealed by allele specific expression"

### Supplementary figure 1

a

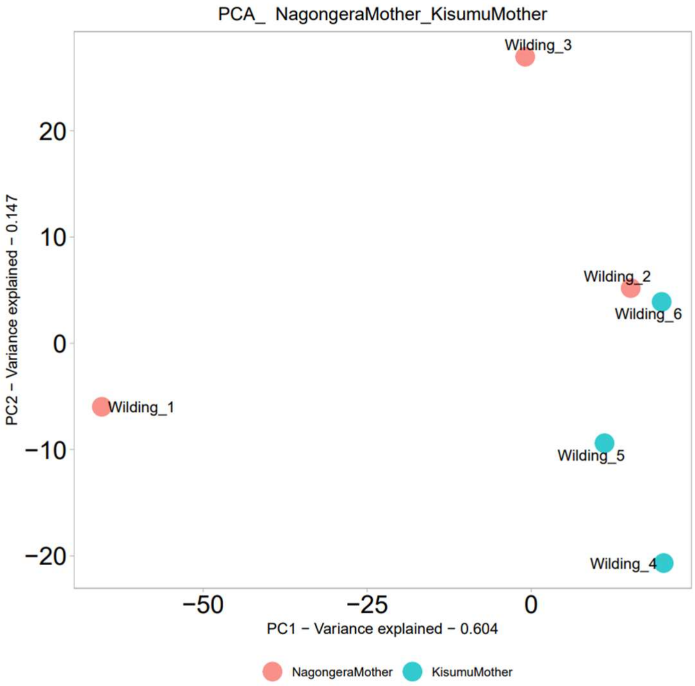


b.

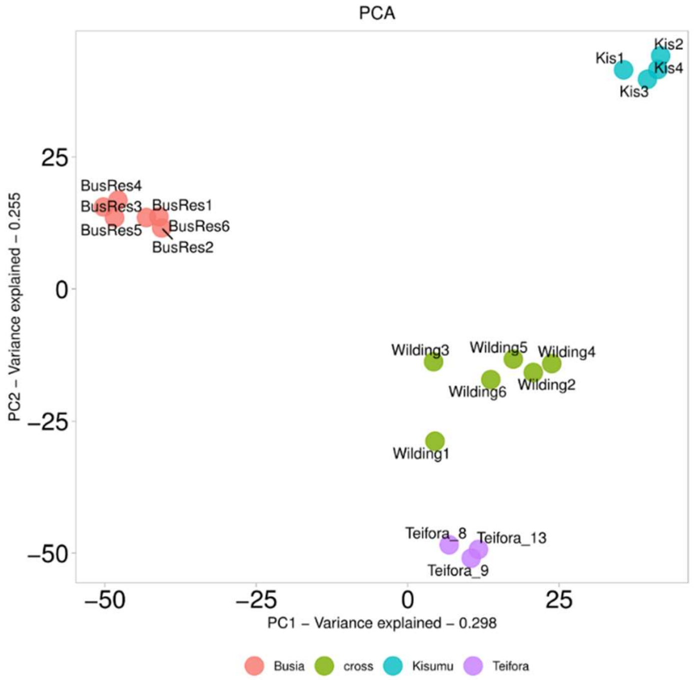


a. PCA of gene expression for reciprocal crosses. Mapping of RNAseq sample names to crosses: Wilding_1: B1, Wilding_2: B3, Wilding_3: B5, Wilding:4: K2, Wilding_5: K4, Wilding_6: K6

b. PCA of gene expression for crosses compared to other *Anopheles gambiae* sl strains: Teifora_9, Teifora_8 and Teifora_13 are Tiefora colony pools, Kis1, Kis2, Kis3 and Kis4 are Kisumu colony pools, BusRes1, BusRes2, BusRes3, BusRes4, BusRes5, BusRes6 are Busia G28 deltamethrin selected colony pools.

**Supplementary figure 2**

a.

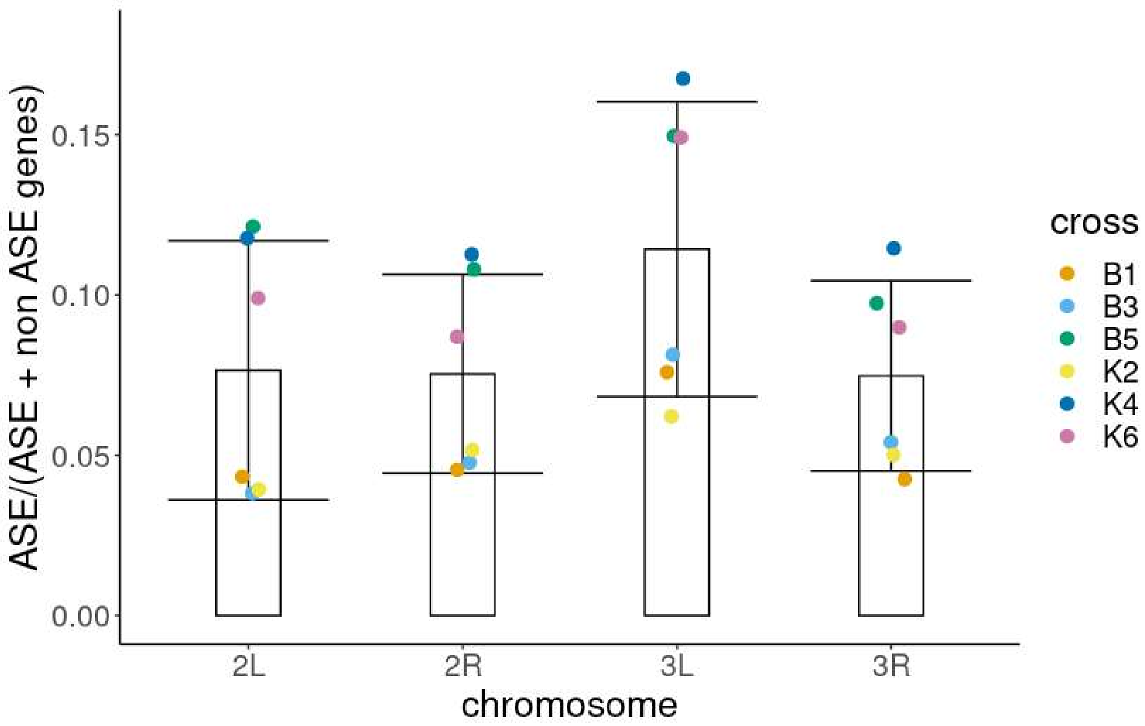


b.

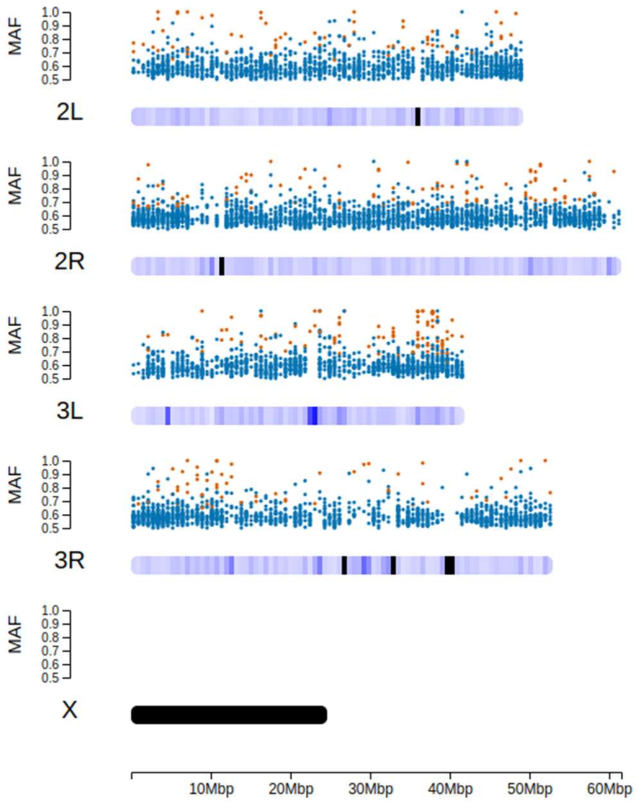

c.

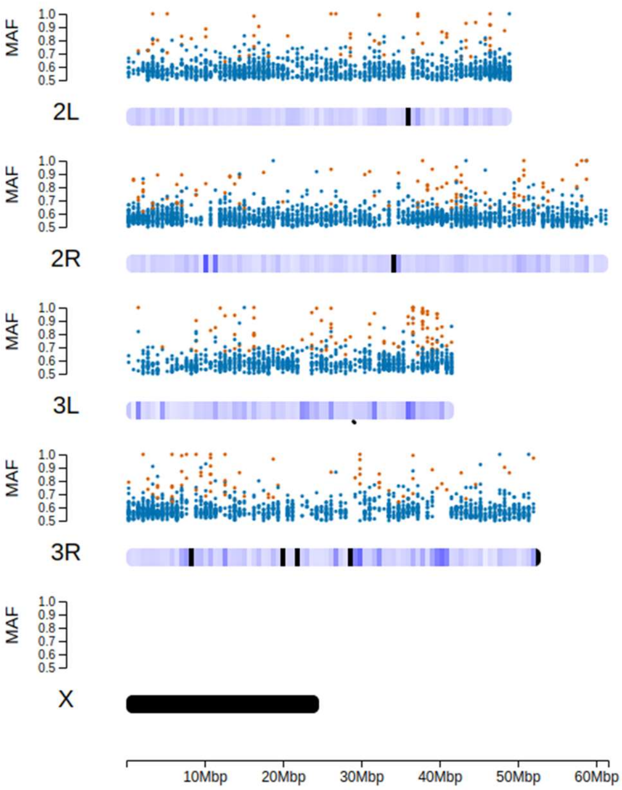
MAF
1.0
0.9
0.8
0.7
0.6
0.5
2L
2R
3L
3R
X
10Mbp
20Mbp
30Mbp
40Mbp
50Mbp
60Mbp

d

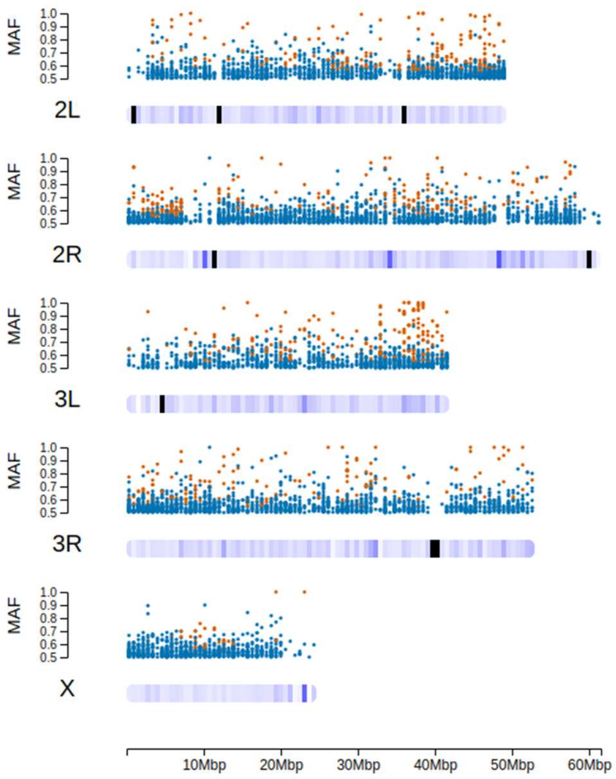
MAF
1.0
0.9
0.8
0.7
0.6
0.5
2L
2R
3L
3R
X
10Mbp
20Mbp
30Mbp
40Mbp
50Mbp
60Mbp

e

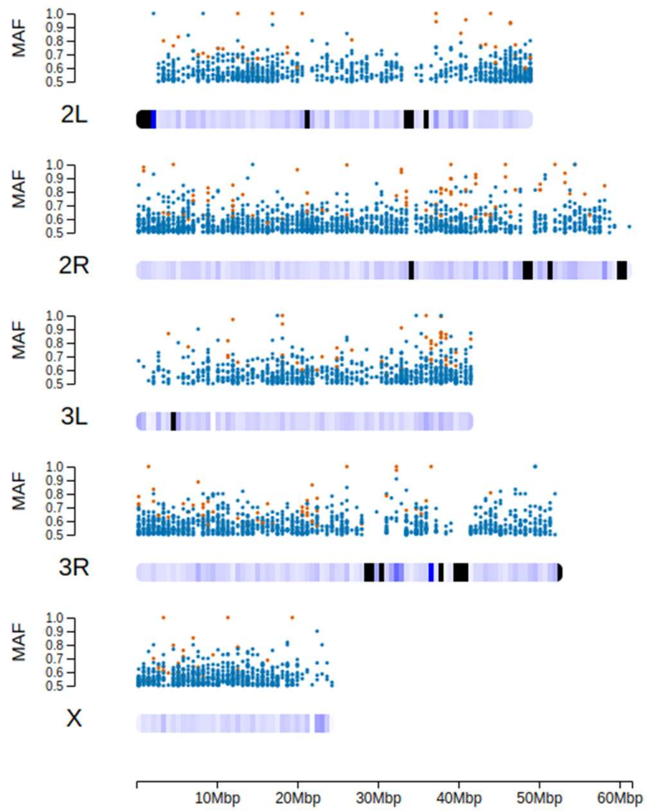
MAF
1.0
0.9
0.8
0.7
0.6
0.5
2L
2R
3L
3R
X
10Mbp
20Mbp
30Mbp
40Mbp
50Mbp
60Mbp

f

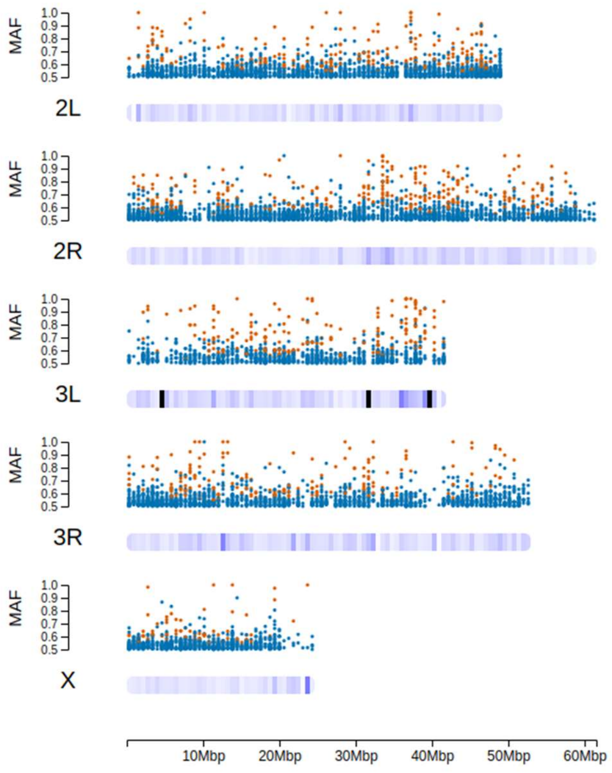
MAF
1.0
0.9
0.8
0.7
0.6
0.5
2L
2R
3L
3R
X
10Mbp
20Mbp
30Mbp
40Mbp
50Mbp
60Mbp

g

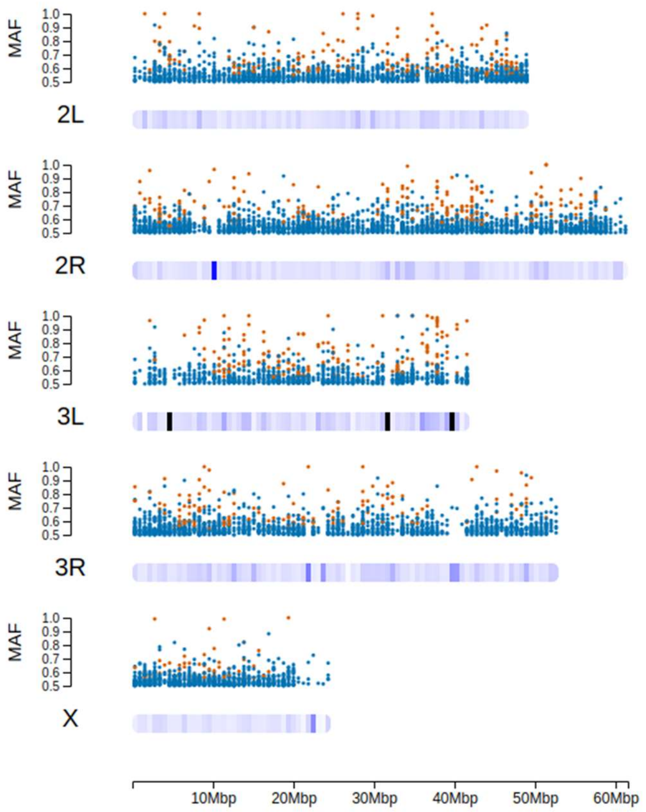


a. Proportion of genes showing allelic imbalance compared to all genes where SNPs were present so ASE could have been detected along each chromosomal arm. Error bars show standard deviation across the 6 crosses.
b. -g. plots of major allele frequency (MAF) along each chromosomal arm. Scatter plot above each chromosomal arm of genes for which the P value for ASE was non-significant (blue), and genes where the P value for ASE was significant at alpha =0.05 following FDR correction (orange). The density of data along the chromosomal arms is on a black to blue to white scale: black regions indicate no data. b. Genes showing ASE and not showing ASE along each chromosome arm for cross B1)
c. Genes showing ASE and not showing ASE along each chromosome arm for cross B3
d. Genes showing ASE and not showing ASE along each chromosome arm for cross B5

e. Genes showing ASE and not showing ASE along each chromosome arm for cross K2

f. Genes showing ASE and not showing ASE along each chromosome arm for cross K4

g. Genes showing ASE and not showing ASE along each chromosome arm for cross K6

### Supplementary Figure 3

Allele balance calculated using SNPs from non-parents

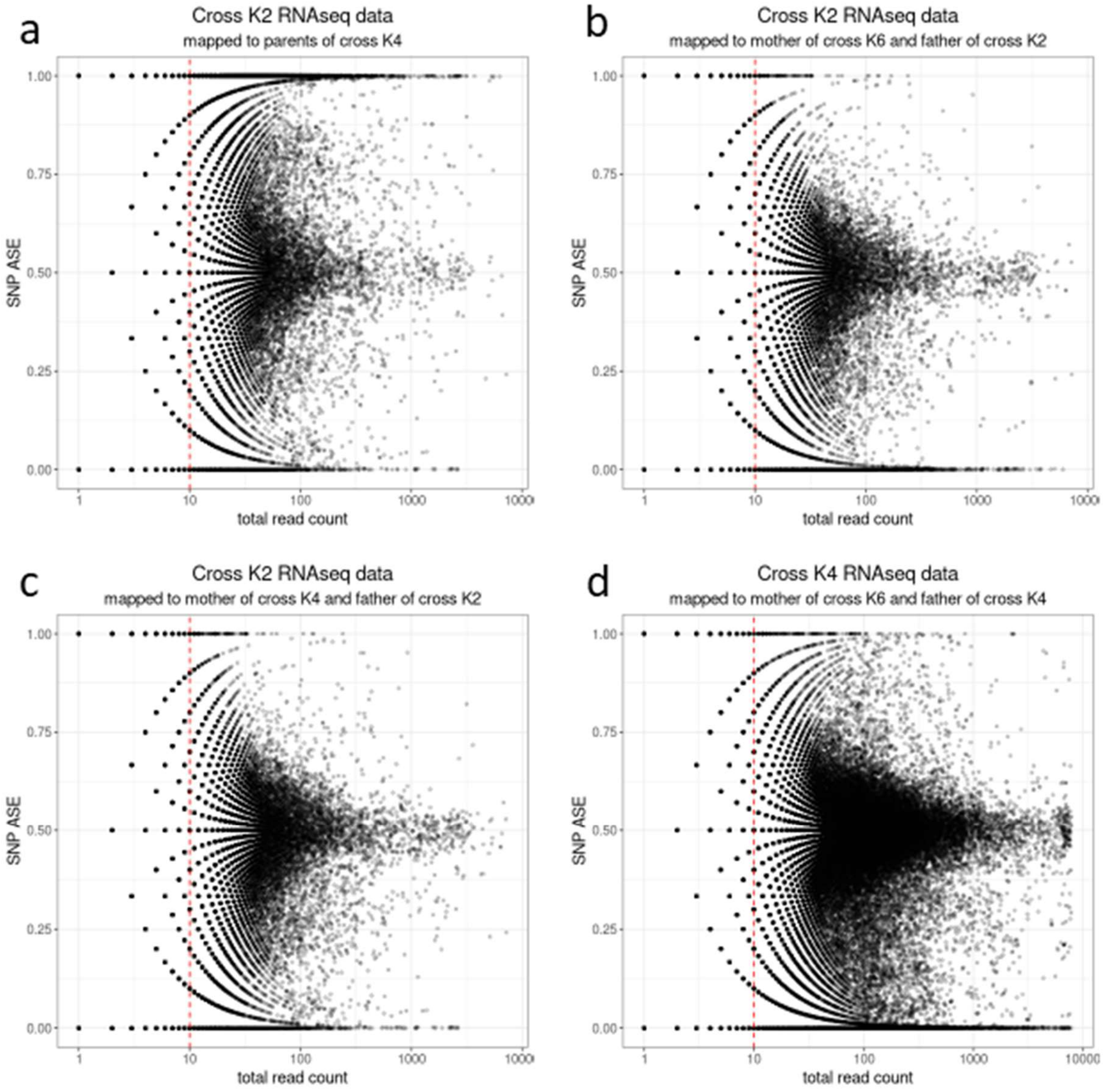


a. RNAseq reads from progeny of cross K2 mapped to K4 parents

b. RNAseq reads from progeny of cross K2 mapped to mother of cross K6 and correct father

c. RNAseq reads from progeny of cross K2 mapped to mother of cross K4 and correct father

d. RNAseq reads from progeny of cross K4 mapped to mother of cross K6 and correct father

### Supplementary Figure 4

Progeny of crosses K4 (Wilding 5) and K6 (Wilding 6) are genetically similar

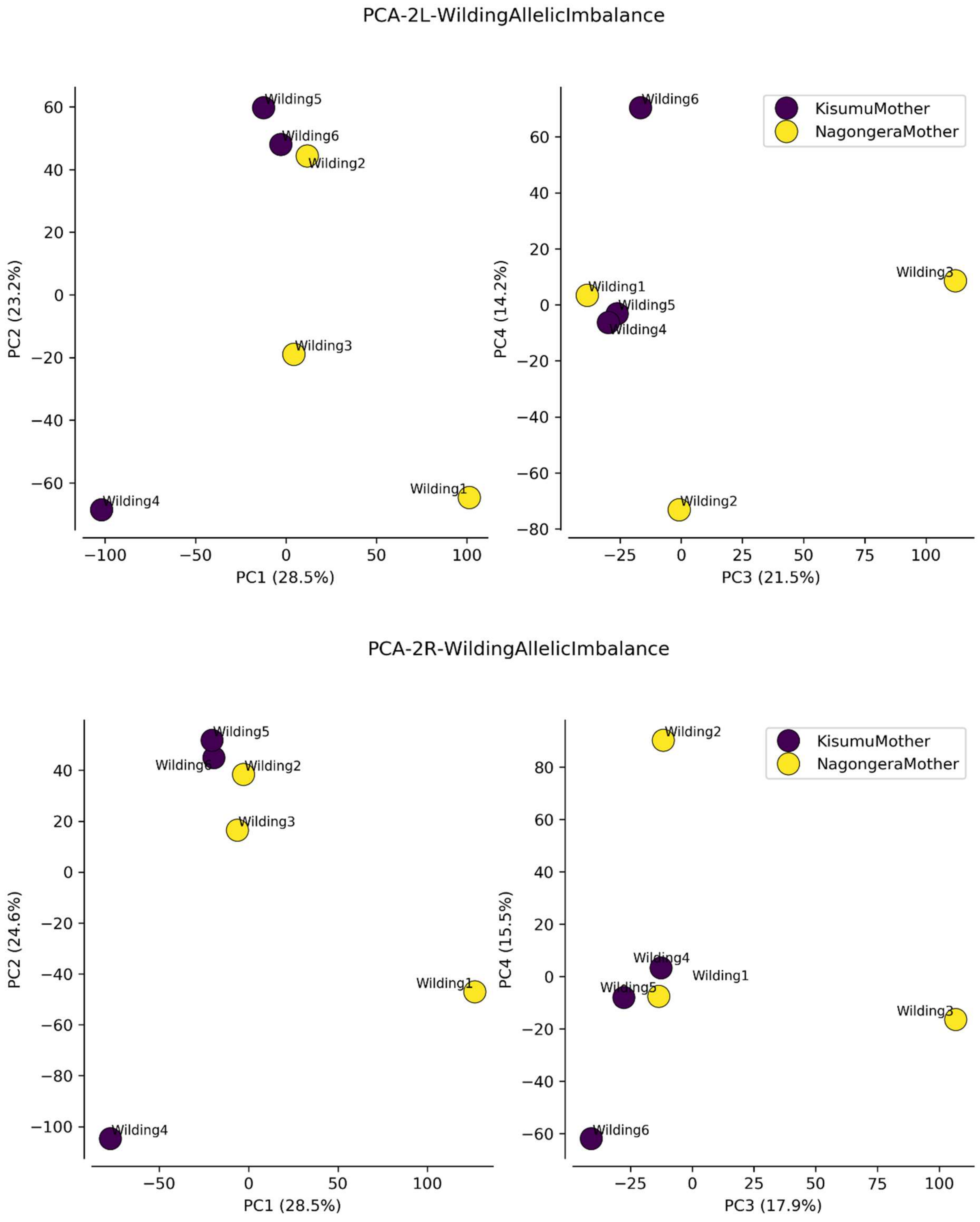

PCA-3L-WildingAllelicImbalance

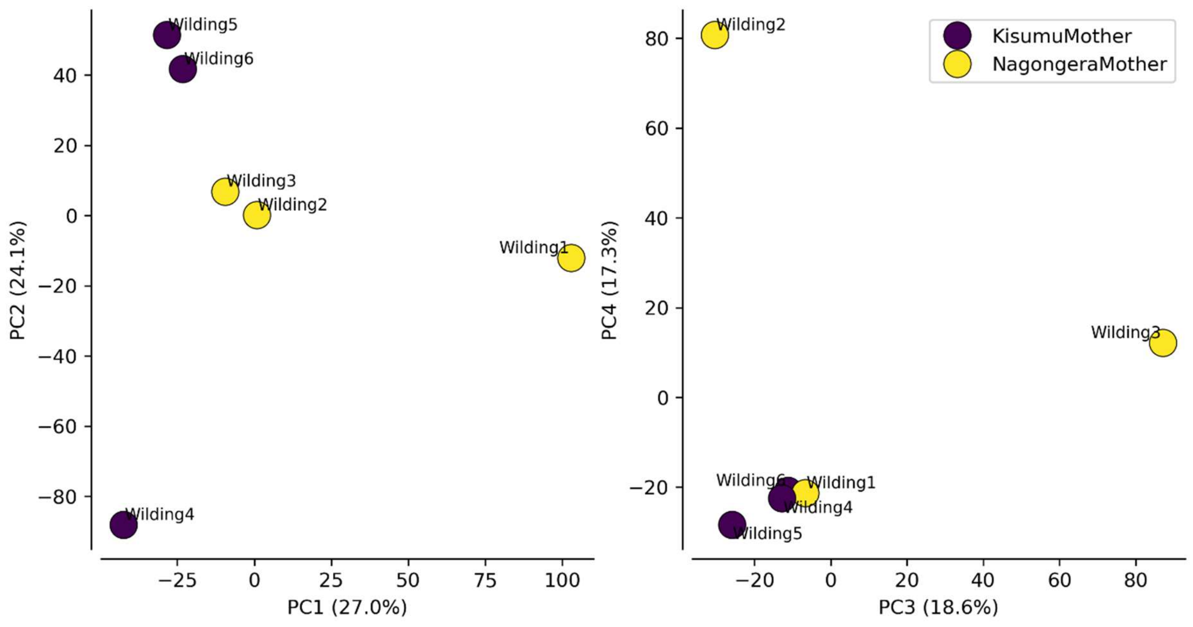


PCA-3R-WildingAllelicImbalance

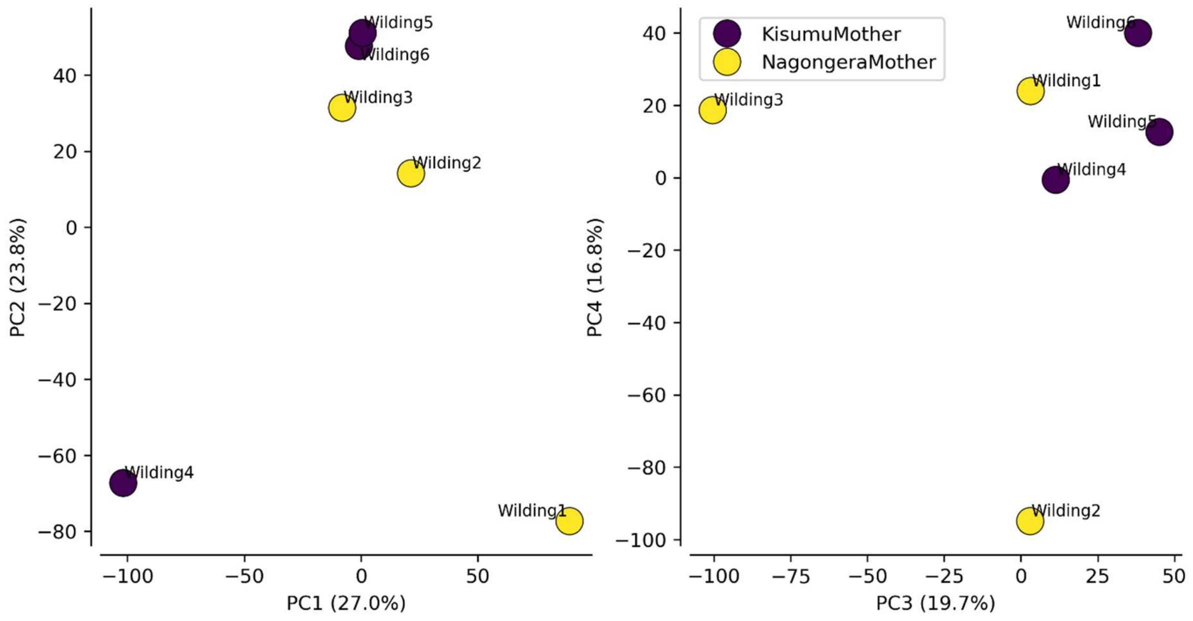

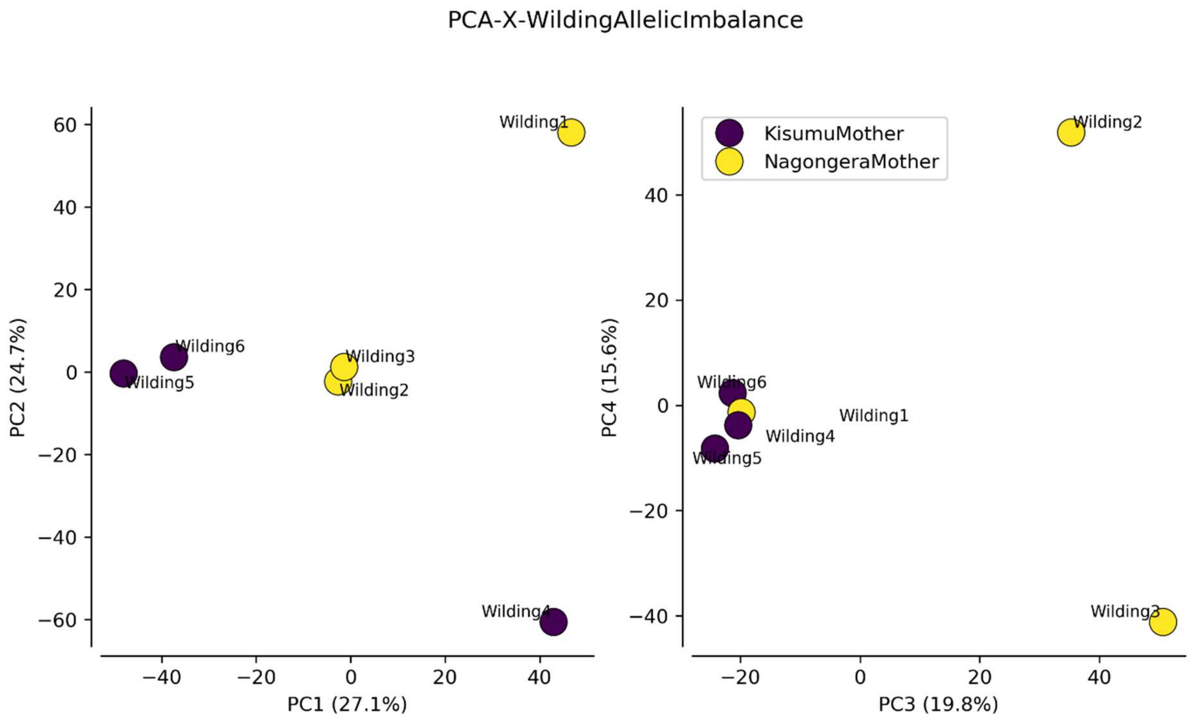


Legend: PCA of sequences inferred by calling SNPs from RNAseq data of crosses between Kisumu and Nagongera colonies. Left plots are, PC1 vs PC2, right plots are PC2 vs PC3. The chromosomal arm for each pair of plots is indicated in the plot titles. % of variance explained by each component is given in brackets in the axis labels.
